# Transcriptome-proteome profiling in *Burkholderia thailandensis* during the transition from exponential to stationary phase

**DOI:** 10.1101/2025.02.17.638721

**Authors:** Ahmed Al-Tohamy, Fabrizio Donnarumma, Anne Grove

## Abstract

Bacterial cells commonly exist in stationary phase, for instance within a host cell. *Burkholderia thailandensis* is closely related to, and a surrogate for highly pathogenic *Burkholderia* species. Understanding the molecular mechanisms that characterize the transition of *B. thailandensis* from exponential to stationary phases is critical to understanding responses to stress or nutrient limitation. We present here an integrated transcriptomic and proteomic analysis of gene and protein expression changes during entry into stationary phase. We identified 928 differentially expressed genes and 832 differentially expressed proteins. Genes encoding proteins involved in benzoate degradation and O-antigen nucleotide sugar biosynthesis were among the most highly upregulated in stationary phase, whereas processes such as translation and flagellar biosynthesis were downregulated. Proteins related to fatty acid degradation and butanoate metabolism accumulated along with proteins involved in synthesis of secondary metabolites. Markedly downregulated proteins included ribosomal proteins as well as the house-keeping iron-sulfur biogenesis proteins. An only modest correlation between transcriptome and proteome changes was seen, and the RpoS sigma factor was not significantly increased during early stationary phase; RpoS is typically abundant during stationary phase and critical for expression of stress-response genes. Our data therefore point to distinct adaptive mechanisms in *B. thailandensis*, including post-translational regulation.

## Introduction

*Burkholderia* is a diverse genus of Gram-negative bacteria, and many species exhibit pathogenicity in plants and animals.^1^ Several of these microorganisms are well-known for their antibiotic resistance and ability to cause severe illness, particularly in immunocompromised individuals such as cystic fibrosis patients.^2^ Other *Burkholderia* species hold biotechnological potential due to their capacity to degrade environmental pollutants.^3^ This dual nature has sparked a growing interest in understanding their metabolic capabilities and antibiotic resistance mechanisms. *Burkholderia thailandensis*, although typically considered non-infectious to humans, serves as an excellent surrogate due to its close evolutionary relationship to pathogenic *Burkholderia* species, particularly *Burkholderia pseudomallei*, which causes melioidosis.^4^

Understanding the metabolic and regulatory pathway dynamics during different growth phases is crucial for understanding bacterial physiology. In nature, the bacteria may experience nutrient-replete environments in which they can grow and divide (feast). However, they are even more likely to encounter nutrient-depleted or stressful conditions under which mere survival becomes paramount (famine). Such adverse conditions, which may for instance be encountered in a host environment, require the bacteria to adjust their gene expression, metabolic, or developmental programs. In the laboratory, these conditions may be simulated by the transition from exponential growth to stationary phase.^5^ During an initial lag phase, which is characterized by metabolic activity without an increase in cell number, the bacteria adapt to their new environment, synthesizing essential proteins and metabolites required for subsequent growth. During the active growth phase, cells divide by binary fission, leading to a rapid increase in population. This phase is marked by high metabolic activity; the population size doubles at regular intervals, resulting in exponential growth.^5^

During the stationary phase, the growth rate slows down due to nutrient depletion and the accumulation of waste products, with no net increase in population density.^5^ However, cells remain metabolically active and undergo significant physiological changes to adapt to the nutrient-limited environment. This phase is distinguished by extensive metabolic reprogramming, including the downregulation of protein synthesis, repression of aerobic metabolism, and the upregulation of alternative energy production pathways.^6^ Entry into stationary phase is a carefully regulated process; one of the early events in Gram-negative bacteria is considered to be accumulation of the alternative sigma factor RpoS, which in turn governs the expression of genes encoding proteins required for survival under suboptimal conditions. Upregulation of RpoS has been reported to be a consequence of changes in transcription, translation, and protein stability.^7^

Amino acid starvation and other nutrient limitation also triggers the stringent response, which is characterized by accumulation of the hyperphosphorylated guanine nucleotides (p)ppGpp. Their synthesis is, for example, triggered by the presence of uncharged tRNAs in the ribosomal A site.^8^ (p)ppGpp then binds the RNA polymerase to affect promoter selectivity and to favor association of RpoS in preference to the housekeeping sigma factor, σ^70^. As a result, rRNA synthesis, ribosomal protein production, and DNA replication are downregulated, while amino acid biosynthesis is upregulated.^9^ Increased utilization of long-chain fatty acids as a carbon source is also characteristic of stationary phase.^10^

Some bacteria also increase production of secondary metabolites upon entry into stationary phase. These compounds can serve various functions, including acting as antibiotics, siderophores, or signaling molecules.^11^ This shift towards secondary metabolism may reflect a competitive strategy as nutrients become scarce.

Regulation of gene expression features prominently as the bacteria adjust to changing environments. Bulk RNA sequencing (RNA-seq) has been extensively used to map such transcriptome changes. However, it is becoming increasingly apparent that post-transcriptional regulation is equally important during the entry into the stationary phase. For example, in *Rhodobacter sphaeroides*, proteome changes often outpace and sometimes diverge from transcriptome changes as the bacteria enter the stationary phase.^12^

Here, we present an integrated view of transcriptome and proteome changes during the transition from exponential to stationary phases of *B. thailandensis*. These insights provide a foundational understanding of the adaptive mechanisms employed by *B. thailandensis* in response to stress or nutrient limitation.

## Experimental Procedures

### Bacterial Strains and Media

The *Burkholderia thailandensis* E264 strain, obtained from the American Type Culture Collection (ATCC), was cultivated in 2×YT medium (16 g/L tryptone, 10 g/L yeast extract, 5 g/L NaCl; adjusted to pH 7.0). Cultures were incubated at 37°C with continuous shaking at 250 rpm. An overnight culture was diluted 1:100 in fresh 2×YT medium. The absorbance at 600 nm was recorded every 2 hours for approximately 24 hours in three biological replicates, with each absorbance representing an average of three technical replicates. Samples representing exponential phase cells were collected at an OD_600_ of 0.6 ± 0.05. The stationary phase cells were collected at an OD_600_ of 2.6 ± 0.05.

### RNA Sequencing (RNA-Seq)

#### Sample Preparation and RNA Extraction

For RNA sequencing, three biological replicates from both the exponential and stationary phases were used. Two mL of culture was collected, followed by centrifugation at 16,000 × g for 2 minutes. The cell pellets were washed twice with diethyl pyrocarbonate (DEPC)-treated water, and then immediately frozen at −80°C. RNA extraction was conducted using the Monarch Total RNA Miniprep Kit (New England BioLabs), following the manufacturer’s guidelines. The integrity of the extracted RNA was assessed by agarose gel electrophoresis. RNA was quantified using a NanoDrop™ 2000c spectrophotometer. An Agilent 2100 Bioanalyzer was used to obtain the RNA Integrity Number (RIN), selecting only samples with a RIN of 8 or higher for further processing.

#### Library preparation and High-Throughput sequencing

Library preparation and high-throughput sequencing was performed by Novogene (https://www.novogene.com). Library preparation was done using the Illumina TruSeq Stranded mRNA Library Prep Kit, adhering to the manufacturer’s instructions. Ribosomal RNAs were depleted from the total RNA, followed by ethanol precipitation. The RNA was then fragmented, and the first strand of cDNA synthesized using random hexamer primers. In the synthesis of the second strand of cDNA, dUTPs were replaced with dTTPs in the reaction buffer. The library was further processed through end repair, A-tailing, adapter ligation, and size selection. Uracil-Specific Excision Reagent (USER) enzyme digestion was applied, followed by amplification and purification. Post-preparation, the libraries were quantified using Qubit and real-time PCR, and a bioanalyzer was used for determination of size distribution. Libraries were pooled, and high-throughput sequencing was executed utilizing a paired-end read configuration on the Illumina NextSeq 500 platform, and Fastq files were generated.

#### Differential gene expression bioinformatics data analysis

The bioinformatics pipeline started with a quality assessment using FastQC.^13^ Post-evaluation, the Trim Galore toolkit^14^ was applied to remove sequencing adapters and unidentified bases (N) and to trim bases of low quality (phred score cutoff = 20). MultiQC was used to assess the quality of the processed dataset.^15^ For alignment purposes, the reference genome of *B. thailandensis* was retrieved from the *Burkholderia* Genome Database (https://www.burkholderia.com).^16^ The genome was indexed using SAMtools. Subsequent alignment of the high-quality reads to this reference genome was performed using the HISAT2 aligner.^17^ Post-alignment, SAMtools was utilized to construct SAM and BAM files from the aligned reads that used for further analysis. Gene expression quantification was conducted using FeatureCounts.^18^ The differentially expressed genes between exponential and stationary phases were identified using DESeq2^19^ based on the count matrix obtained. Genes were considered significantly differentially expressed if they exhibited a log2-fold change of ≥2 or ≤-2 and if they met the statistical significance threshold with a false discovery rate (FDR) of <0.05. This FDR value was adjusted for multiple testing using the Benjamini-Hochberg procedure.^20^

#### Data visualization

Heatmaps were constructed using the ComplexHeatmap package in R.^21^ Log2-fold change values were converted to z-scores, normalizing gene expression across samples. Hierarchical clustering was applied to group genes with similar expression patterns. To identify and visualize differentially expressed genes, Volcano plots were generated using the Plotly Express library in Python.^22^

### Quantitative Mass Spectrometry Proteomics

#### Protein extraction, Tandem Mass Tag (TMT) labeling and fractionation

We followed the filter-aided sample preparation (FASP) method^23^ with the following modifications. Forty mL of the bacterial culture were centrifuged at 17,800 × g for 5 min, followed by two washes with phosphate buffered saline (PBS). The bacterial pellets were stored at −80°C. Protein extraction was carried out using a lysis buffer (8 M urea in 50 mM Tris, pH 8.0). The cell pellets were sonicated twice using a Branson SFX250 Sonifier at 35% output at 10 secs pulse with 1-minute gap in between. Homogenized samples were centrifuged at 8,600 × g for 5 min. Protein concentration was determined using the Pierce™ bicinchoninic acid (BCA) Protein Assay Kit (Thermo Fisher Scientific). Proteins (100 µg total) were first reduced with 50 mM DTT and then alkylated with iodoacetamide. Subsequent overnight digestion was performed with trypsin at 37°C. TMTsixplex™ labeling was conducted following the manufacturer’s protocol (Thermo Fisher Scientific). Following labeling, the peptides were pooled and fractionated using strong cation exchange stage tips (AttractSPE Disk, Affinisep, Le Houlme, Normandy, France). A total of 6 fractions where dried and resuspended in 0.1% formic acid. The LC-MS/MS analysis was executed on an Orbitrap Fusion Tribrid mass spectrometer. Detailed procedures may be found in the Supporting Information.

#### Data acquisition

Mass spectrometry analysis was conducted on a Q-Exactive orbitrap mass spectrometer (Thermo Scientific, Waltham, MA) coupled to an Ultimate 3000 RSLC liquid chromatography system (Thermo Scientific). A binary mobile phase system was employed both for sample loading and trapping, as well as for analytical separation. Sample injection was set to 5 µL, and each fraction was resuspended in 10 µL of 0.1% formic acid. The mobile phase composition was as follows: A = LC-MS grade H_2_O with 0.1% formic acid; B = 100% acetonitrile with 0.1% formic acid. The sample was loaded by connecting the capillary carrying the tryptic peptides to an Acclaim PepMap 100 C18 trap cartridge (0.3 x 5 mm, 5 µm particle size, 100 Å pore size) and transferred using a mobile phase composition of 2% B over the course of 5 minutes at a flow rate of 20 µL/min. Separation was carried on an Acclaim PepMap 100 C18 nanoLC column (0.075 × 250 mm, 3 µm particle size, 100 Å pore size) using a linear gradient from 6% to 35% mobile phase B over the course of 120 minutes at a flow rate of 300 nL/min. The mass spectrometer was operated in data-dependent acquisition with a loop count of 10. Full MS parameters were as follows: resolution = 70,000, automatic gain control (AGC) = 1 × 10^6^, scan range = 300-1600 m/z and maximum injection time = 30 ms. The data-dependent acquisition parameters were as follows: resolution = 35,000, AGC target = 5 × 10^4^, isolation window = 2 m/z, maximum injection time = 50 ms and normalized collision energy = 30. Additional settings included: dynamic exclusion = 60 seconds, peptide match = preferred, excluded isotopes = on, charge exclusion = unassigned +7, +8 and >+8 charges, and maximum AGC target = 500, which resulted in an intensity threshold of 10,000.

#### Proteomics Data Analysis

The proteomics experiments yielded raw mass spectra, which were subjected to data processing using Proteome Discoverer software.^24^ This encompassed peak picking, spectral alignment, and peptide identification. For peptide identification, the SEQUEST HT search engine was used. This engine matched the acquired spectra against the *Burkholderia thailandensis* protein database, which was obtained from UniProt.^25^ Quantification of protein abundances relied on the measurement of TMT reporter ion intensities. Proteins were considered significantly differentially expressed if they exhibited a log2-fold change of ≥ 0.5 or ≤ −0.5, and if they met the statistical significance threshold with a P-value of < 0.05.

#### Data Visualization

Principal Component Analysis (PCA) was conducted using the PCA function from the sklearn.decomposition module in Python.^26^ The analysis was performed on standardized proteomic data to reduce dimensionality and highlight the variance between the experimental conditions. The PCA results were visualized using matplotlib.^27^ The Volcano plot was generated as described above.

### Protein-Protein interaction (PPI) network construction

To construct the protein-protein interaction (PPI) network for *B. thailandensis*, the relevant protein FASTA files were retrieved from the UniProt database.^25^ Since *B. thailandensis* is not available in the STRING available organisms,^28^ we generated a custom reference proteomic dataset using the STRING server based on this FASTA data. The differentially upregulated proteins were then used to construct the PPI network.

PPI data was sourced from the STRING database. The dataset included interactions with combined confidence scores, reflecting the likelihood of functional associations between protein pairs. Further analysis of the PPI network was performed using custom Python script using NetworkX.^29^ We calculated each protein number of interactions, average interaction score, degree centrality, betweenness centrality, closeness centrality, eigenvector centrality, clustering coefficient, interacting proteins list, cluster numbers, and average shortest path length. In the PPI, nodes represent proteins, and edges represent interactions between proteins. To identify functional modules within the PPI network, the Louvain method for community detection was employed.^30^

### KEGG Pathway changes and Gene Ontology (GO) analysis

Gene Set Enrichment Analysis (GSEA) was performed to identify significantly regulated pathways using the KEGG database.^31^ Pathways were identified using the fgsea package in R.^32^ Pathway enrichment results were visualized using custom scripts to generate bubble plots, displaying the normalized enrichment score (NES), -log10(p-value), and counts of genes/proteins.

Selected pathways were further visualized using the Pathview package in R.^32^ DEGs/DEPs, represented as log2-fold change (log2FC) values, were mapped onto KEGG pathways. Custom color schemes were applied, with upregulated genes shown in red and downregulated genes in blue, where the intensity of the color correlates with the magnitude of expression changes.

GO Enrichment Analysis was conducted to identify overrepresented GO terms within DEGs and DEPs. GO terms were categorized into Biological Process (BP), Molecular Function (MF), and Cellular Component (CC) categories. The analysis was performed using custom Python and R scripts to assess the enrichment of GO terms. The enrichment results were filtered to include only those terms with a p-value < 0.05. The enriched GO terms for each category were visualized using bar plots generated with ggplot2,^33^ where the x-axis represents the count of genes, and the y-axis represents the GO terms with the color scheme reflecting the level of statistical significance. Specifically, more intense red colors represent higher p-values, indicating lower statistical significance, while more intense blue colors represent lower p-values, indicating higher significance.

### Integration of RNA-seq and proteomics data

To integrate the RNA-Seq and proteomics data, gene-protein mapping was performed using curated annotations from the UniProt database. The DEGs and DEPs were merged based on their respective gene or protein identifiers using a custom Python script. The integrated dataset was then analyzed to identify genes and proteins with opposite or consistent regulation between RNA-Seq and proteomics data.

Pearson correlation^34^ analysis was performed to assess the relationship between RNA-Seq and proteomics data. Log2-fold changes from both datasets were normalized using the StandardScaler function from the sklearn library.^26^ Correlation coefficients and p-values were calculated using the scipy library ^35^. Scatter plots with regression line were generated to visualize correlations, and density plots were created to compare the distributions of normalized log2-fold changes. Violin plots were used to compare the distribution of normalized log2-fold changes between the RNA-Seq and proteomics datasets. For visualization, seaborn ^36^ and matplotlib^27^ libraries were used.

### RpoS Sigma Factor Analysis

#### RT-qPCR

An overnight culture of *B. thailandensis* was diluted 1:100 and sub-cultured in 2×YT medium, and samples were collected at various OD_600_ values (0.1, 0.2, 0.4, 0.6, 1.1, 1.4, 1.6, 1.9, 2.2, 2.4, 2.6, 2.8, and 3.0). Two mL of the cell culture was pelleted, washed twice with autoclaved DEPC-treated water, and stored at −80°C. RNA was extracted utilizing the Monarch total RNA miniprep kit (New England Biolabs, Ipswich, MA) according to the manufacturer’s protocol. RNA was electrophoresed on agarose gels to ascertain integrity. Quantitative RT-PCR was performed using Luna one-step universal master mix (New England Biolabs). Data represent means (±SDs) from biological triplicates (each determined from technical triplicates) using the comparative threshold cycle (*C_T_*) method (2^−ΔΔ*CT*^) for which *hgprt* (*BTH_I1148*) was used as a reference gene. The *C_T_* values for *hgprt* were constant under the conditions used for these experiments. The primers sequences can be found in Supplemental Table S9.

#### Comparison of regulons

The reported RpoS regulons from *B. pseudomallei* and *E. coli* were compared. Both reported regulons were based on 2D gel electrophoresis followed by protein identification by mass spectrometry. For *B. pseudomallei*, 58 unique differentially expressed proteins were identified.^37^ The RpoS regulon of *E. coli* included 35 proteins.^38^ BlastP searches were used to identify homologous proteins in *B. thailandensis.*^39^

## Results

### Transcriptome Profiling

The growth of *B. thailandensis* was monitored, and cells representing exponential growth (OD_600_∼0.6) and stationary phase (OD_600_∼2.6) were collected for transcriptome profiling (Figure 1A). For each condition, cells were collected from three independent cultures (three biological replicates). Differentially expressed genes (DEGs) exhibiting a significant log2-fold change greater than |2| were primarily examined to evaluate transcriptome changes occurring on entry into stationary phase. Based on this cutoff, we identified 928 DEGs, of which 564 were upregulated and 364 were downregulated (Figure 1B). The top 20 up- or downregulated genes are shown in Table 1; the entire dataset is included in Supplemental Table S1. Hierarchical cluster analysis of DEGs visualized using a heatmap reflects the clustering of samples with similar expression patterns (Figure 1C).

**Figure 1.**
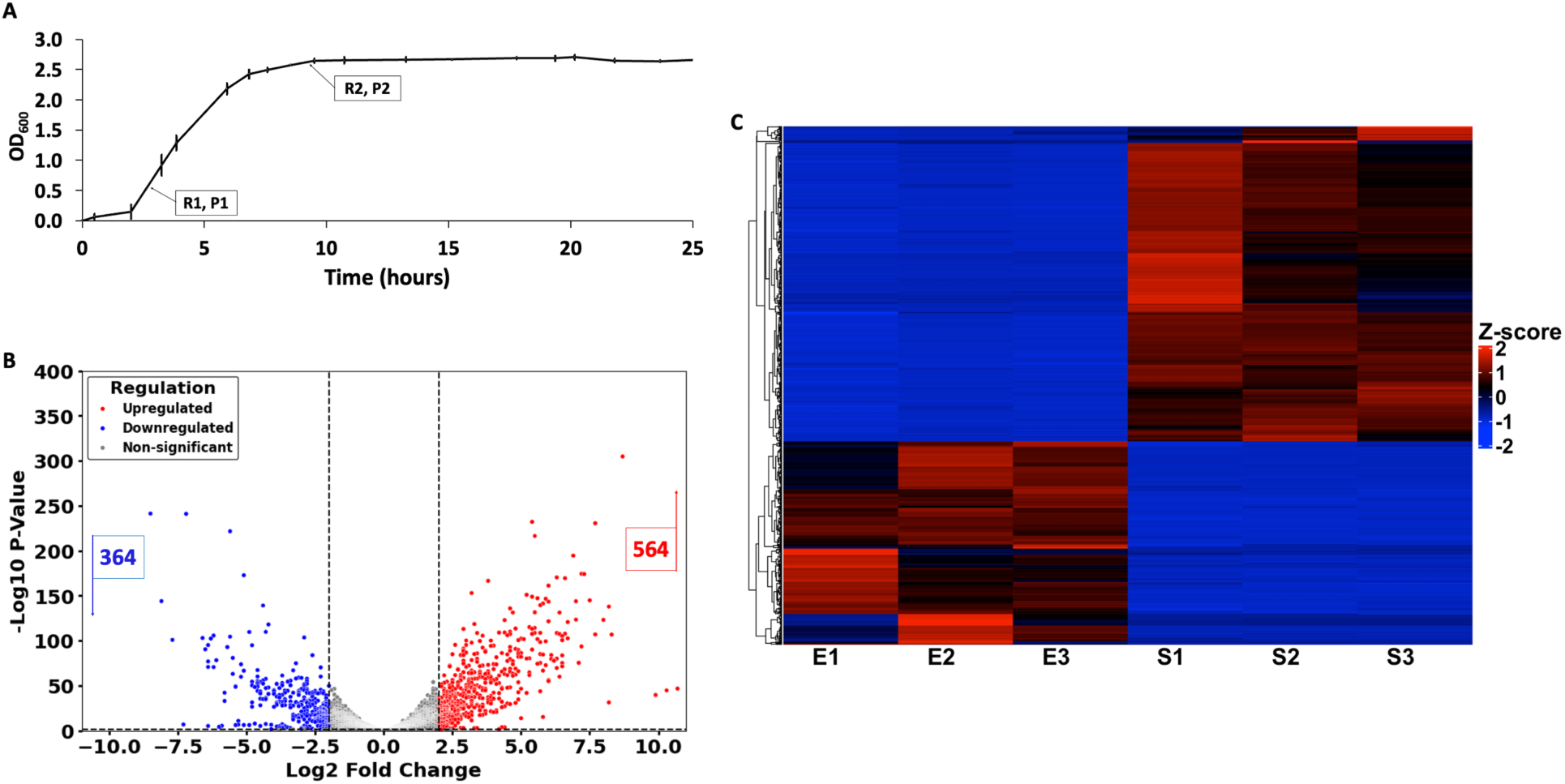
Growth curve and gene expression analysis. A. Growth curve of *B. thailandensis*: lag phase (OD_600_ ∼ 0.0-0.2), exponential growth (OD_600_ ∼ 0.2-2.4), and stationary phase (OD_600_ ∼ 2.5- 2.7). Error bars represent standard deviation from three biological replicates. Labels R1 and P1 indicate the point where cells were collected for RNA and protein isolation during exponential phase, while R2 and P2 indicate the corresponding point for the stationary phase. B. Volcano plot of RNA-seq data. Significantly up-regulated genes (564) in red and down-regulated genes (364) in blue. The horizontal dashed line hugging the x-axis represents a p-value of 0.05, while the vertical dashed lines indicate log2-fold changes of |2|. C. Heatmap of gene expression changes in exponential and stationary phase. Red indicates higher expression and blue indicates lower expression. Hierarchical clustering on both gene and sample axes is included to display patterns of gene expression co-regulation. Samples E1, E2, and E3 represent three biological replicates for the exponential phase, while samples S1, S2, and S3 represent three biological replicates for the stationary phase.

**Table 1.**
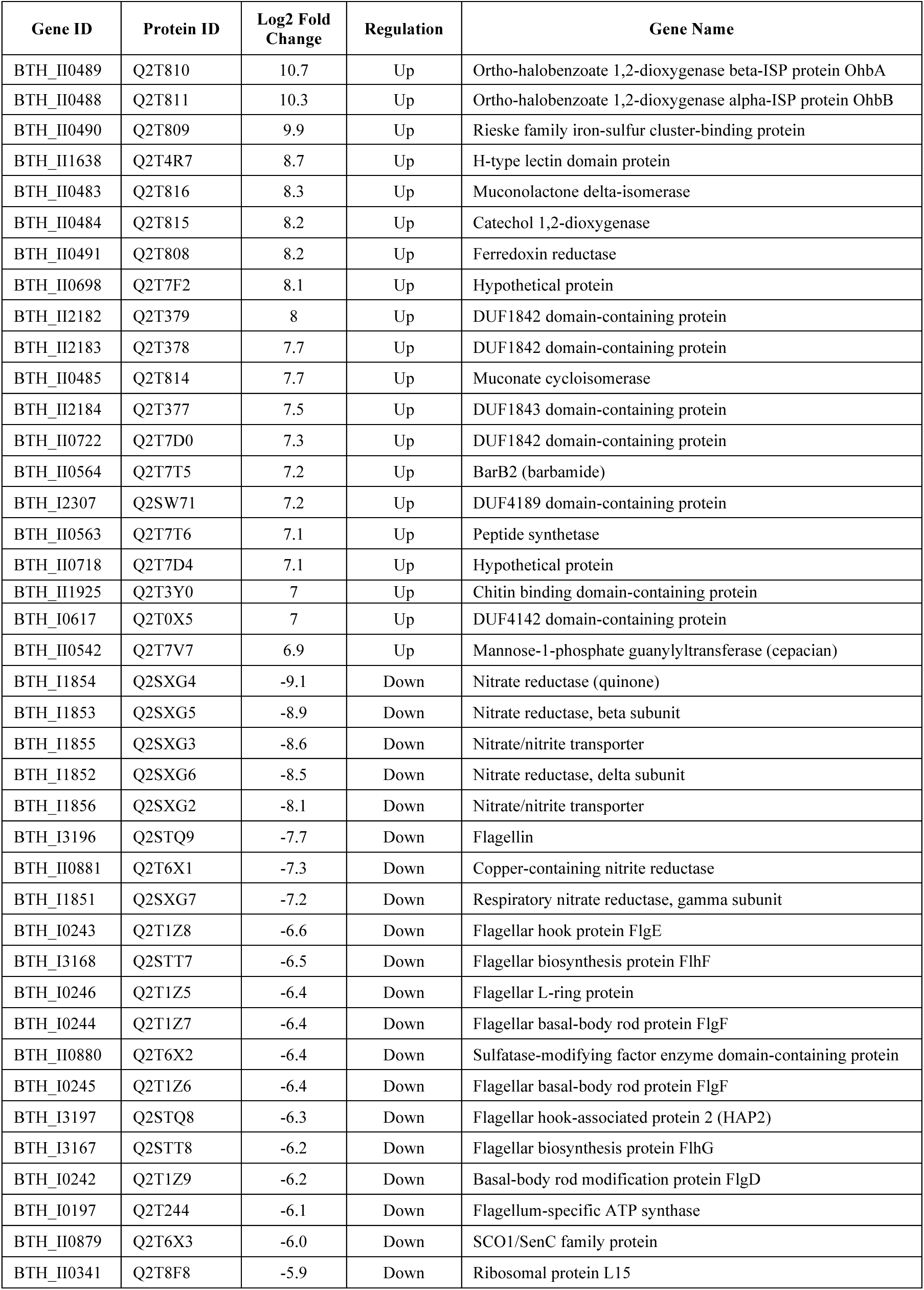
Top Differentially Expressed Genes.

To evaluate the biological functions of the DEGs, they were functionally categorized using the Kyoto Encyclopedia of Genes and Genomes (KEGG) pathway classification (Figure 2A and Supplemental Table S2). The top pathways involved in bacterial adaptation to stationary phase were selected. Significantly downregulated pathways included nitrogen metabolism, chemotaxis, and flagellar assembly, and genes encoding two-component systems were also generally downregulated. Upregulated pathways included benzoate degradation and biosynthesis of nucleotide sugars.

**Figure 2.**
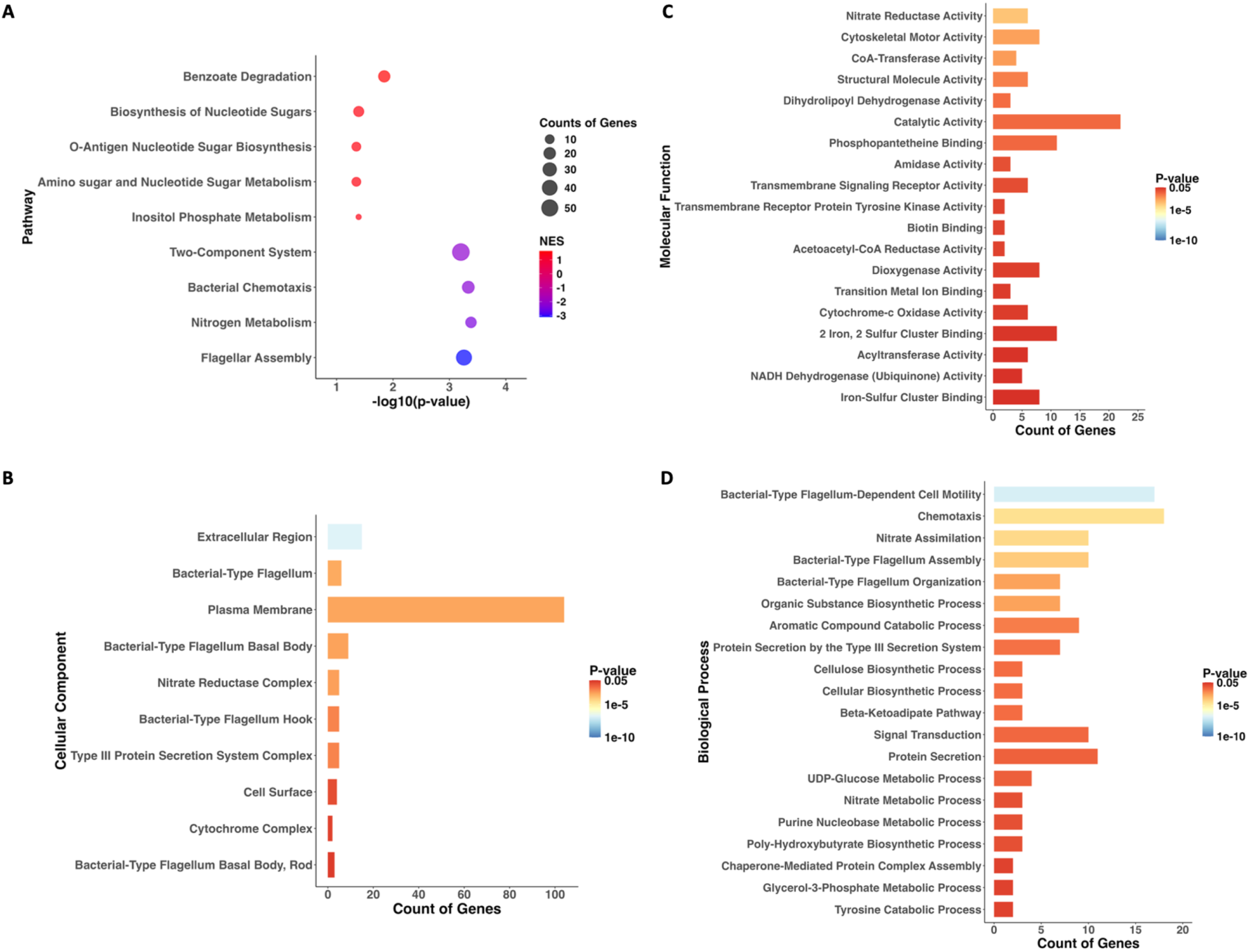
RNA sequencing KEGG pathways and gene ontology. A. Bubble plot illustrating significantly regulated KEGG pathways during the stationary phase compared to the exponential phase. Each pathway is represented by a circle, with the size of the circle indicating the count of the DEGs in the pathway. The x-axis represents the -log10(p-value) of the change. The color of the circles indicates the Normalized Enrichment Score (NES), with red shades representing upregulated pathways (NES closer to 1) and blue shades representing downregulated pathways (NES closer to −3). B.-D. Bar plots showing the enriched Gene Ontology (GO) terms among differentially expressed genes for Cellular Component, Molecular Function and Biological Process, respectively. The x-axes show the observed gene count associated with each GO term. The color of each bar corresponds to the calculated p-value, with more red colors representing higher levels of enrichment significance (lower p-value) and more blue colors representing lower levels of enrichment significance (higher p-value).

Gene Ontology (GO) annotations were calculated to provide a framework for describing altered functions. In the Cellular Component category, by far the largest category was plasma membrane, while categories also represented in the KEGG pathway analysis were also represented, including bacterial-type flagellum and nitrate reductase complex (Figure 2B). The Molecular Function category revealed alterations in catalytic activity as well as categories related to iron-sulfur cluster metabolism (Figure 2C). In the Biological Process category, several terms related to metabolic processes such as nitrate assimilation and aromatic compound cCatabolic process were changed (Figure 2D).

### Downregulated Pathways Based on Transcriptome Profiling

Chemotaxis and flagellar assembly were significantly downregulated (Table 1 and Supplemental Figure S1). This includes genes encoding proteins such as methyl-accepting chemotaxis proteins (MCPs), CheA, and CheY, which are involved in chemotactic signal transduction. CheA is part of a two-component system that initiates the chemotactic signal transduction cascade.^40^ The flagellar assembly pathway includes genes such as *fliC*, which encodes flagellin, a crucial element for flagella formation and function. Also downregulated was *fliA* (log2-fold change −1.7), which encodes the flagellar sigma factor, a member of the σ^28^ protein family, which directs RNA polymerase to transcribe flagellar genes. Such reduction in motility is characteristic of bacteria as they transition from the motile, planktonic state to a more sedentary lifestyle as the resource demand to fuel motility becomes too high.^41^

*Escherichia coli* encodes three nitrate reductases, of which one is periplasmic and two are membrane-bound. One operon encoding a membrane-bound nitrate reductase is under control of the two-component system NarXL, which responds to nitrate and nitrite. The other is under control of RpoS, and its expression is increased in stationary phase.^42^ Such redundancy is also seen in *B. thailandensis.* We found that *BTH_I1852-1854*, which is downstream from genes encoding NarXL was among the most downregulated genes (Table 1 and Supplemental Figure S2). By contrast, *BTH_II1249-1251*, encoding the other membrane-bound nitrate reductase is upregulated, perhaps mediated by RpoS.

In addition, we observed the expected downregulation of genes encoding ribosomal proteins (log2-fold range −5.9 to −1.5; Supplemental Table S1). Genes encoding proteins that participate in the arginine deiminase pathway (*arcDABC; BTH_I1283-1286*) were also markedly downregulated. This pathway is involved in ATP generation during anaerobic growth.^43^ Also notable was the downregulation of genes encoding Type III Secretion System components and effectors, genes encoding enzymes involved in synthesis and export of the antibacterial 4-hydroxy-3-methyl-2-alkenylquinolines (HMAQ), NADH dehydrogenase, involved in electron transport, and the *suf* operon, encoding proteins involved in iron-sulfur cluster biogenesis under iron limitation or oxidative stress.^44^

### Upregulated Pathways Based on Transcriptome Profiling

Benzoate degradation was markedly upregulated in stationary phase. Benzoate may derive from degradation of aromatic compounds, and in some species phenylalanine, and the steps involved in its aerobic degradation include its conversion to catechol and further processing in the β-ketoadipate pathway (Supplemental Figure S3).^45^ Upregulated genes include *BTH_II0471-0473*, encoding benzoate dioxygenase catalyzing the first step towards conversion to catechol as well as all genes encoding enzymes required for conversion of catechol to succinyl-CoA (Table 1). Also upregulated was the operon *BTH_II0488-0491* encoding the anthranilate-inducible ortho-halobenzoate dioxygenase, which converts anthranilate to catechol^46^; anthranilate (2- aminobenzoate) is an intermediate in the synthesis and degradation of tryptophan.^47^

Other notable upregulated gene clusters included *BTH_II0562-0574,* encoding proteins with homology to the barbamide biosynthetic enzymes from *Lyngbya majuscule* and *BTH_II0542-0552*, encoding biosynthetic enzymes for the exopolysaccharide cepacian. Biosynthesis of nucleotide sugars were also upregulated, reflecting a need for building blocks of carbohydrates and glycoconjugates.

### Proteome Profiling

Changes in protein levels were monitored using quantitative mass spectrometry, using the same samples used for transcriptomics, resulting in identification of 3,033 proteins of a total of 5,760 (for details on the workflow, see Supplemental Figure S4). Using a log2-fold change cutoff of |0.5|, we identified 832 differentially expressed proteins (DEPs), with 552 upregulated and 280 downregulated during the stationary phase (Figure 3A-B). The principal component analysis (PCA) revealed particularly tight clustering of samples collected in stationary phase (Figure 3C). The top 20 accumulating or depleted proteins are shown in Table 2, and the entire dataset is provided in Supplemental Table S3. Several key pathways were highlighted through KEGG pathway analysis (Figure 4A and Supplemental Table S4). Expected changes included a decrease in ribosome biogenesis and increased accumulation of proteins related to fatty acid degradation, glycolysis, and amino acid metabolism.

**Figure 3.**
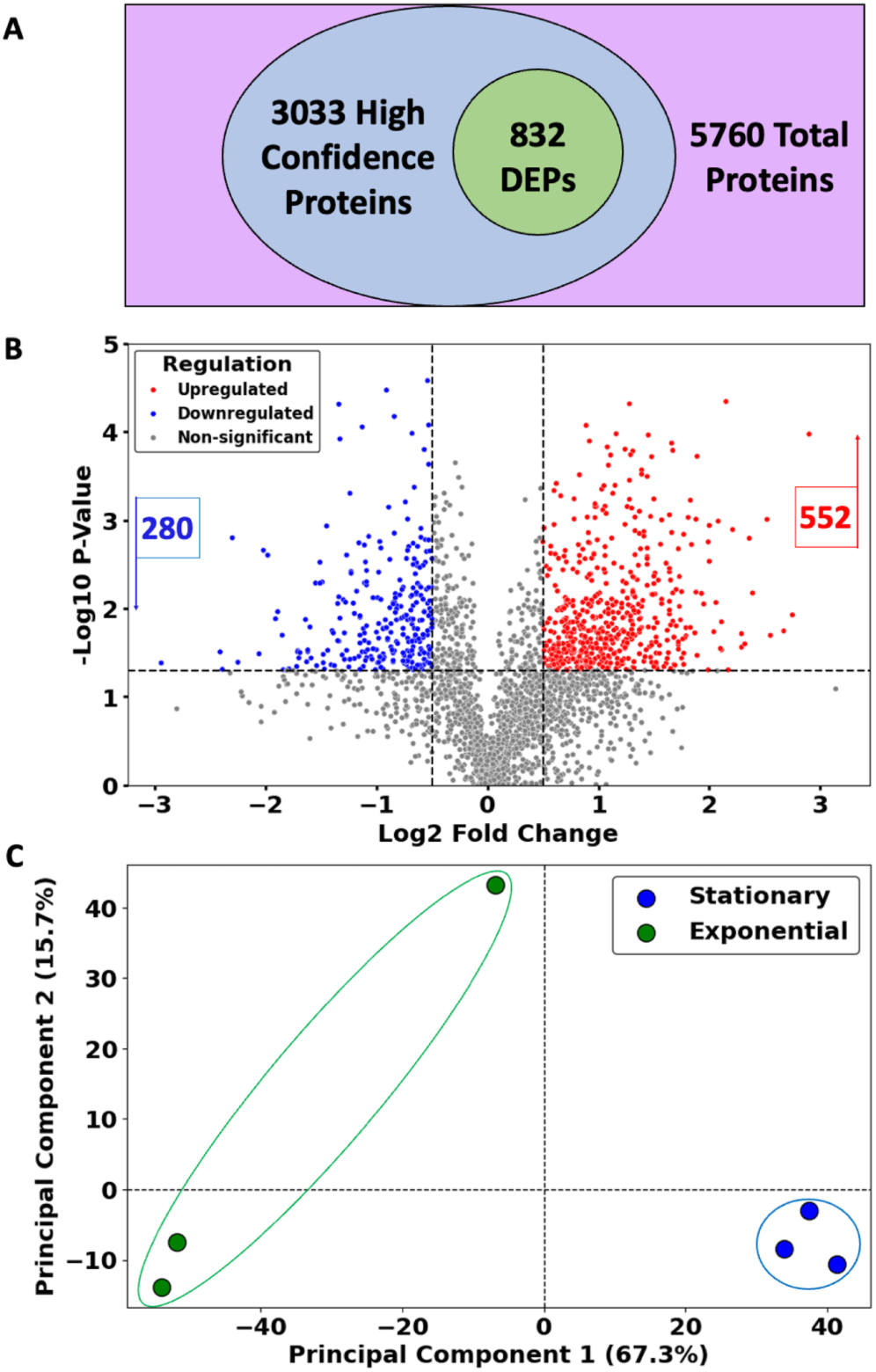
Mass spectrometry quantitative proteomics profiling. A. Venn diagram illustrating the overlap of differentially expressed proteins (832) relative to proteins identified with high confidence (3033). The total number of proteins (5760) in *B. thailandensis* is indicated. B. Volcano plot of differentially expressed proteins in stationary relative to exponential phase. The horizontal dashed line represents a p-value of 0.05 and the vertical dashed lines represent the log2-fold change of |0.5|. Upregulated proteins are shown in red (552), while downregulated proteins are shown in blue (280). C. Principal Component Analysis (PCA) of proteomic profiles. Each point represents a biological replicate, with colors representing the exponential phase (green) and the stationary phase (blue).

**Figure 4.**
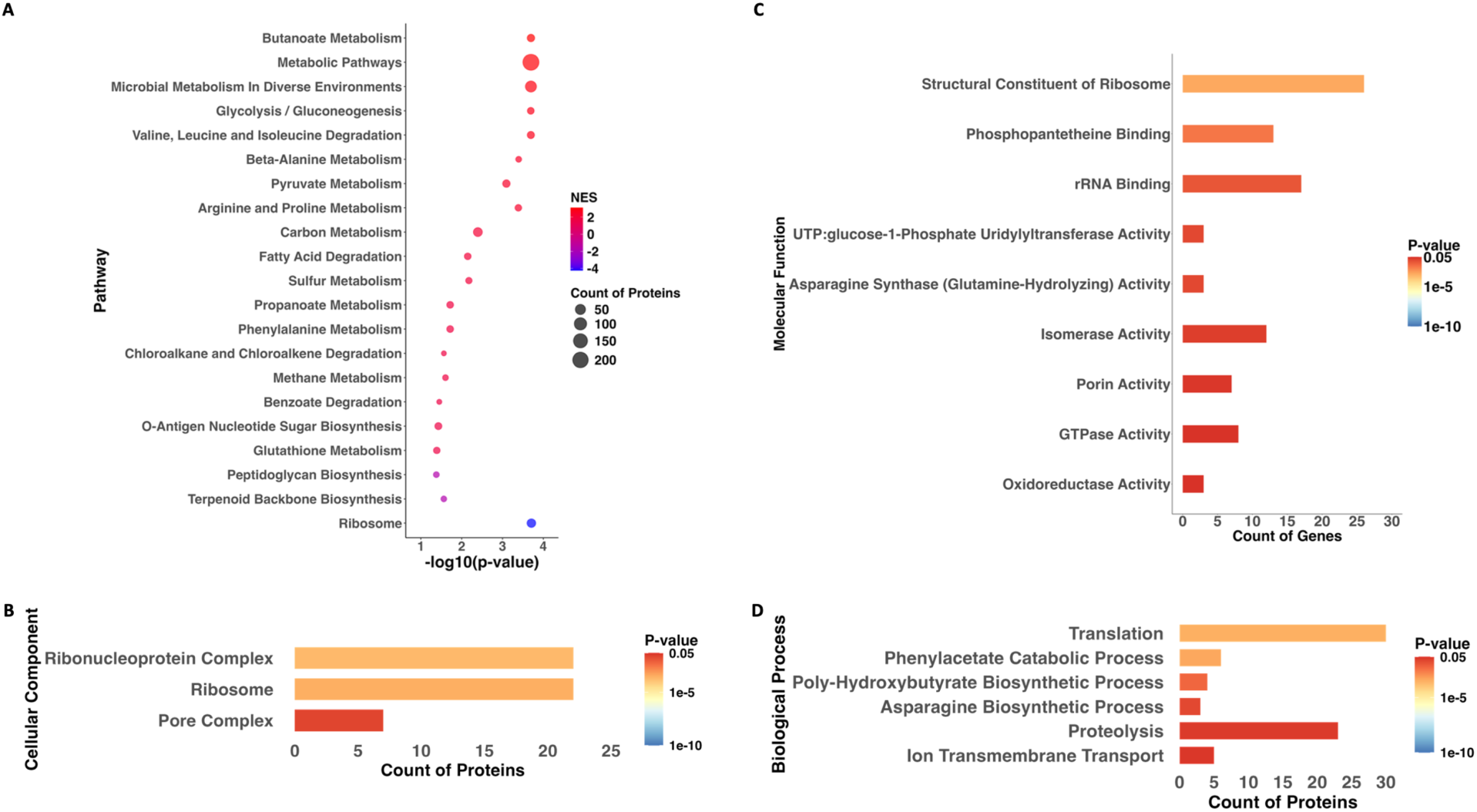
Quantitative proteomics KEGG pathways and gene ontology. A. Bubble plot illustrating the altered KEGG pathways during the stationary phase compared to the exponential phase. Each circle represents a pathway, with the size of the circle indicating the count of the DEPs in the pathway. The x-axis represents the -log10(p-value) of the change. The color of the circles indicates the Normalized Enrichment Score (NES), with red shades representing upregulated pathways (NES closer to 2) and blue shades representing downregulated pathways (NES closer to −4). B-D. Bar plots showing the enriched Gene Ontology (GO) terms among differentially expressed proteins for Cellular Component, Molecular Function and Biological Process, respectively. The x-axis shows the observed protein count associated with each GO term. The color of each bar corresponds to the calculated p-value, with more red colors representing higher levels of enrichment significance (lower p-value) and more blue colors representing lower levels of enrichment significance (higher p-value).

**Table 2.**
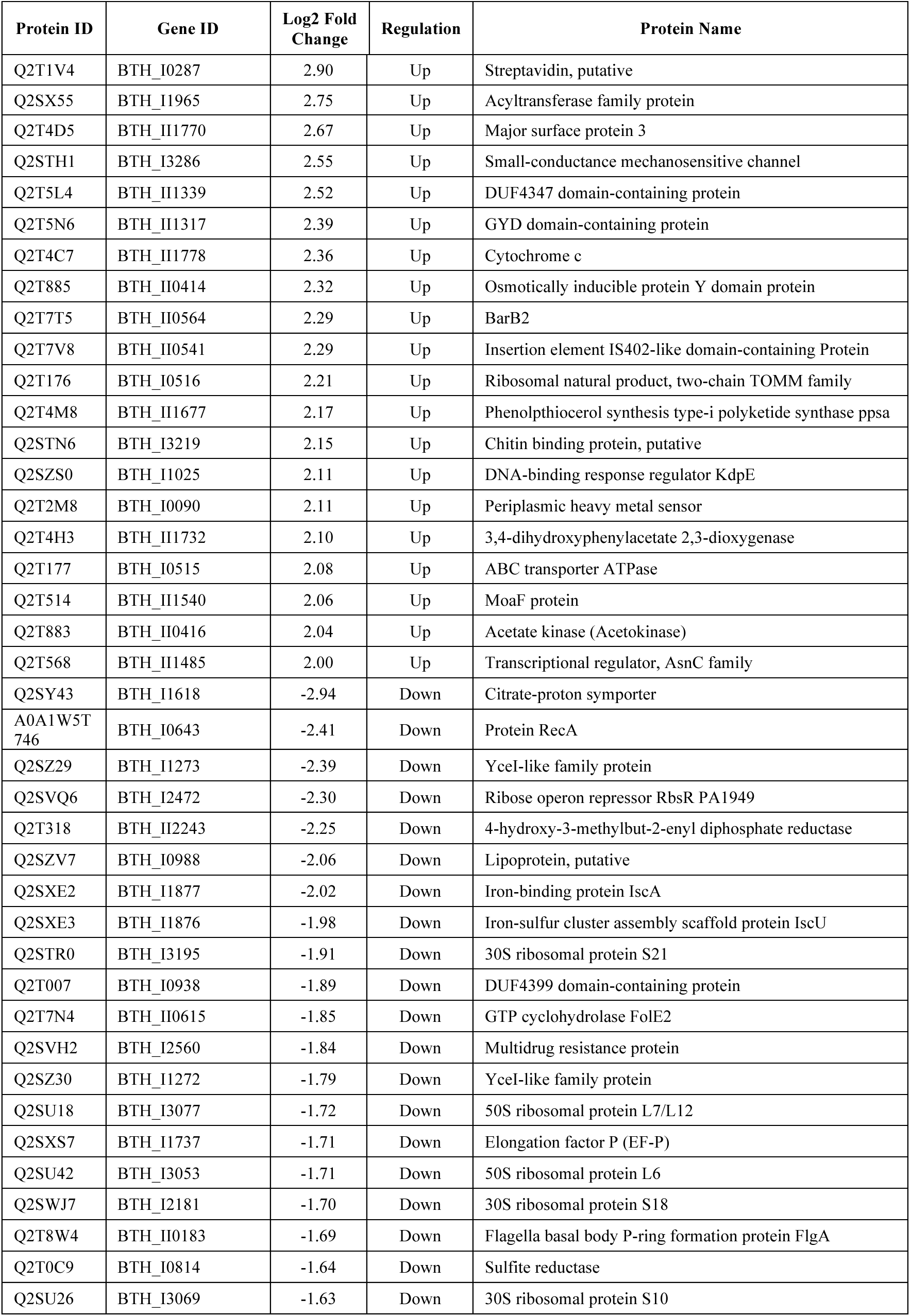
Top Differentially Expressed Proteins.

The GO analysis reflected the expected change in Ribosome and Ribonucleoprotein Complex along with Pore Complex, all in the Cellular Component category (Figure 4B). The Molecular Function category also reflected changes in these categories (structural constituent of ribosome and rRNA binding). Other important molecular functions included oxidoreductase activity and phosphopantetheine binding (Figure 4C). In the Biological Process category, the most altered terms included translation and proteolysis (Figure 4D).

### Downregulated Proteins

The ribosome biogenesis pathway, one of the most energy-consuming biological processes,^9^ was significantly downregulated during the stationary phase (Supplemental Figure S5). Ribosomal proteins were among the downregulated DEPs, along with several proteins required for translation, such as elongation factors G and P and peptide chain release factor 2. FtsZ, required for cell division, was also down, and so was the bifunctional protein PurH, which catalyzes two steps in purine *de novo* biosynthesis. Also down was the peptidoglycan biosynthesis pathway, including proteins such as peptidoglycan D,D-transpeptidase MrdA and penicillin-binding protein, all reflecting the reduced growth and the need to conserve resources. We also noted marked downregulation of the house-keeping iron-sulfur cluster assembly proteins IscS, IscU, and IscA (Table 2).

### Upregulated Proteins

KEGG pathways classified as metabolic pathways and microbial metabolism in diverse environments were among the most markedly upregulated (Figure 4A). Metabolism of fatty acids was increased, as expected. Metabolism of butanoate was significantly upregulated, including upregulation of polyhydroxybutyrate depolymerase; polyhydroxybutyrate serves as a storage form of carbon. In addition, beta-hydroxybutyrate is one of the ketone bodies that accumulate during fatty acid degradation when there is insufficient oxaloacetate for entry of acetyl-CoA into the citric acid cycle (Supplemental Figure S6).^48^ Metabolism of phenylalanine and tryptophan (Figure 4A and Supplemental Table S4) were upregulated; such upregulation is consistent with the observation that genes encoding proteins involved in degradation of benzoate and anthranilate were upregulated.^47^ Proteins involved in biosynthesis of several secondary metabolites were also seen to accumulate, including proteins related to biosynthesis of barbamide, malleilactone, and thailandamide.

The glutathione metabolism pathway was significantly upregulated, with key proteins such as glutathione S-transferase involved in detoxification processes to cope with increased oxidative stress (Figure 4A).^49^ Consistent with an oxidative stress response, we also noted accumulation of alkyl hydroperoxide reductase and *trans*-aconitate methyltransferase; the latter has a role in preventing accumulation of *trans*-aconitate, which is generated when the iron-sulfur cluster of the citric acid cycle enzyme aconitase is damaged by oxidative stress ^50^. DpsA (DNA protection during stress) was also upregulated, as expected (Supplemental Table S2).

The nitrate reductase encoded by *BTH_II1249-1251* was seen to accumulate, consistent with upregulation of the corresponding genes. There was also significant accumulation of a xanthine dehydrogenase subunit; this enzyme functions in purine salvage where it promotes conversion of ATP to GTP. This in turn favors synthesis of the alarmone (p)ppGpp.^51^ Accumulation of a diguanylate cyclase, which generates the second messenger cyclic di-GMP (c- di-GMP), was also noted (Supplemental Table S2). In most bacterial species, including *B. thailandensis*, elevated levels of c-di-GMP are associated with reduced motility and increased biofilm formation.^52^

### Protein-Protein Interaction Networks (PPI) During the Stationary Phase

A predicted protein-protein interaction (PPI) network was constructed, focusing on upregulated proteins identified by mass spectrometry. This approach was employed to pinpoint proteins that may be actively engaged in the bacterial response to stationary phase conditions. The resulting PPI network consisted of 552 nodes (proteins) and 839 edges (interactions), with an average node degree of 3.04, and a maximum degree of 26. Notably, the observed number of edges (839) significantly exceeded the expected number (587), as evidenced by a PPI enrichment p-value of <1.0 × 10^-16^. This enrichment suggests a highly interconnected network, indicative of a robust and coordinated system that surpasses random chance.

Functional enrichment analysis revealed stationary phase PPIs associated with metabolic processes such as fatty acid and amino acid metabolism, reflecting the metabolic reprogramming necessary for survival under nutrient-limited conditions (Figure 5). Corresponding KEGG pathways and GO processes are detailed in Supplemental Figure S7, with topological analysis results of each node provided in Supplemental Table S5.

**Figure 5.**
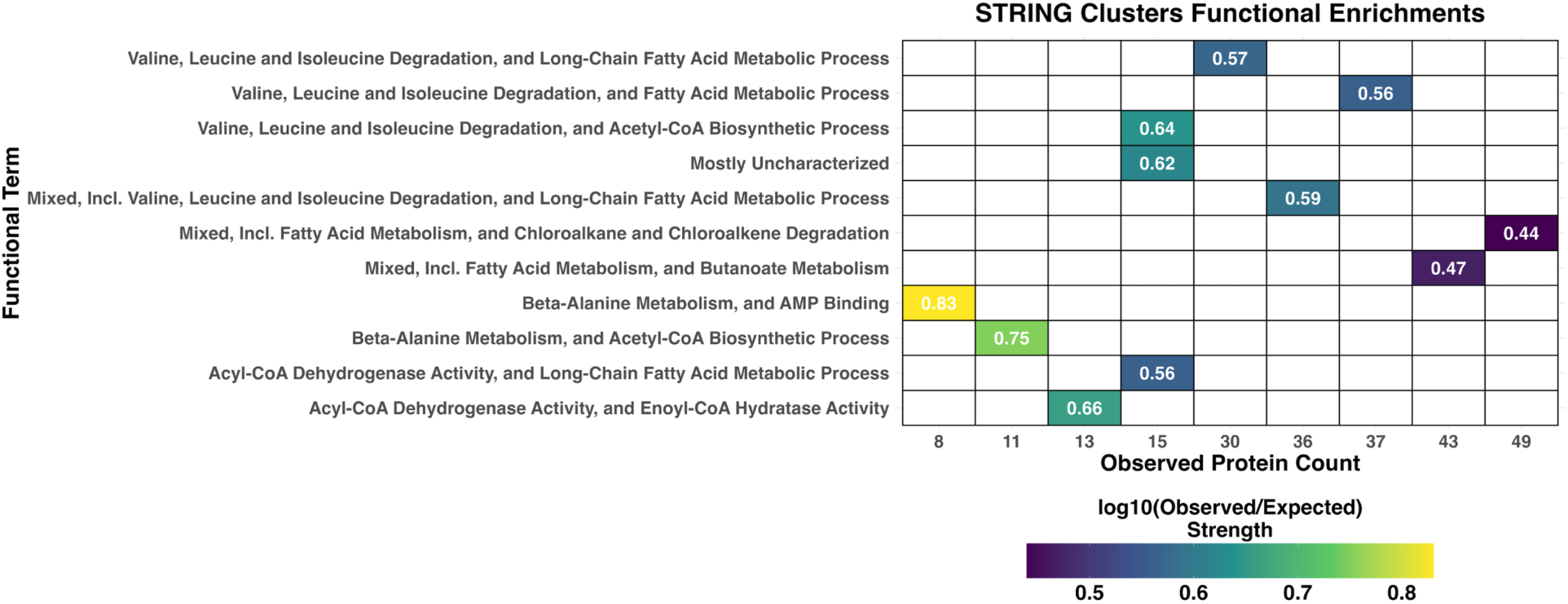
Protein-protein interaction (PPI) network and functional enrichment. Heatmap displaying the enrichment strength of STRING clusters based on the observed gene count for each functional term description. The color intensity represents the log10 of the observed to expected ratio, with darker colors signifying stronger interactions.

Formate Dehydrogenase (FDH) beta subunit (Q2T7E2) was identified as the hub protein with the most (26) interacting partners (Table 3). FDH is integral to the microbial metabolism of one-carbon compounds, particularly within the formate oxidation pathway, which is essential for maintaining redox balance under nutrient-limited conditions.^53^ The interaction of FDH with enzymes such as adenylylsulfate kinase and cysteine synthase suggests a potential linkage between formate metabolism and sulfur assimilation, which is crucial for synthesizing sulfur-containing amino acids and cofactors. Additionally, interactions with malate synthase and phosphoenolpyruvate carboxykinase suggest FDH’s involvement in coordinating the glyoxylate cycle and gluconeogenesis, thereby reorganizing carbon flux during periods of nutrient scarcity. The network further reveals interactions between FDH and nitrate reductase subunits, suggesting its association with the utilization of alternative electron acceptors under the oxygen-limited conditions typical of the stationary phase. Furthermore, interactions with proteins associated with aromatic compound degradation could provide an alternative carbon source under nutrient-limited conditions.

**Table 3.** Main Hub Proteins.

| Hub Protein | Protein Annotation | Number of Interactions |
| --- | --- | --- |
| Q2T7E2 | Formate dehydrogenase, beta subunit | 26 |
| Q2T882 | Phosphate acetyl butyryltransferase family protein | 24 |
| Q2STF6 | Bifunctional protein PutA | 22 |
| Q2T4A5 | 3-hydroxyisobutyrate dehydrogenase (HIBADH) | 22 |
| Q2T066 | Aldehyde dehydrogenase family protein | 21 |
| Q2T2J3 | Acyl-CoA dehydrogenase | 21 |
| Q2SWN6 | Fatty oxidation complex, alpha subunit, putative | 20 |
| Q2T4A2 | Acyl-CoA dehydrogenase | 20 |
| Q2T4A3 | AMP-binding enzyme | 19 |
| Q2SUJ0 | Phenylacetate degradation probable enoyl-CoA hydratase paaB | 17 |
| Q2SWA7 | Aldehyde dehydrogenase | 17 |
| Q2T125 | Enoyl-CoA hydratase/isomerase family protein | 16 |
| Q2T883 | Acetate kinase (acetokinase) | 14 |
| Q2SZL3 | Acyl-CoA dehydrogenase domain protein | 14 |
| Q2T4A4 | Methylmalonate-semialdehyde dehydrogenase (CoA acylating) | 14 |
| Q2T6Q3 | Enoyl-CoA hydratase/isomerase family protein | 14 |
| Q2T3K0 | Malate synthase G | 13 |
| Q2SYR3 | Glutaryl-CoA dehydrogenase (ETF) | 13 |
| Q2STC4 | Methylmalonate-semialdehyde dehydrogenase (CoA acylating) | 13 |
| Q2T1J8 | Fatty oxidation complex, alpha subunit, putative | 13 |
| Q2T4A6 | Enoyl-CoA hydratase/isomerase family protein | 13 |
| Q2SX31 | Malate synthase | 12 |
| Q2SWC1 | Acetoacetyl-CoA reductase | 12 |
| Q2T7X9 | Acyl-CoA dehydrogenase domain protein | 12 |

**Table 3**. The UniProt accession number for the main hub proteins are shown, along with a functional description and the total number of interacting proteins.

Phosphate acetyl/butyryltransferase family protein (Q2T882) was identified as another critical hub within the PPI network, interacting with 25 other proteins. Interacting proteins with roles in fatty acid β-oxidation, branched-chain amino acid metabolism and proteins involved in energy storage and mobilization via polyhydroxyalkanoates (PHAs), which is essential for survival during prolonged stationary phase conditions, speak to the integration of distinct metabolic pathways. Connections with malate synthase and isocitrate lyase further emphasize the rerouting of carbon metabolism through the glyoxylate cycle.

The network analysis generally highlights significant clusters involved in fatty acid metabolism and amino acid degradation, such as the cluster related to valine, leucine, and isoleucine degradation (Figure 5). Taken together, these findings indicate a strategic reallocation of resources to maintain metabolic flexibility and cellular integrity under nutrient-limited conditions.

### Integrative Analysis of RNA-Seq and Quantitative Proteomics Data

To gain a comprehensive understanding of the molecular changes in *B. thailandensis* during the stationary phase, we performed an integrative analysis of RNA-Seq and quantitative proteomics data. This analysis aimed to correlate mRNA levels with protein abundance and to identify shared and divergent regulatory mechanisms at the transcriptional and translational levels. We first assessed the correlation between RNA-Seq and proteomics data by comparing the normalized log2-fold changes of differentially expressed genes and proteins. A scatter plot was used to visualize this relationship (Figure 6A), with a Pearson correlation coefficient (r = 0.4) indicating a moderate positive correlation. This moderate correlation is consistent with previous studies, which have shown that mRNA and protein levels do not always correlate strongly.^54^

**Figure 6.**
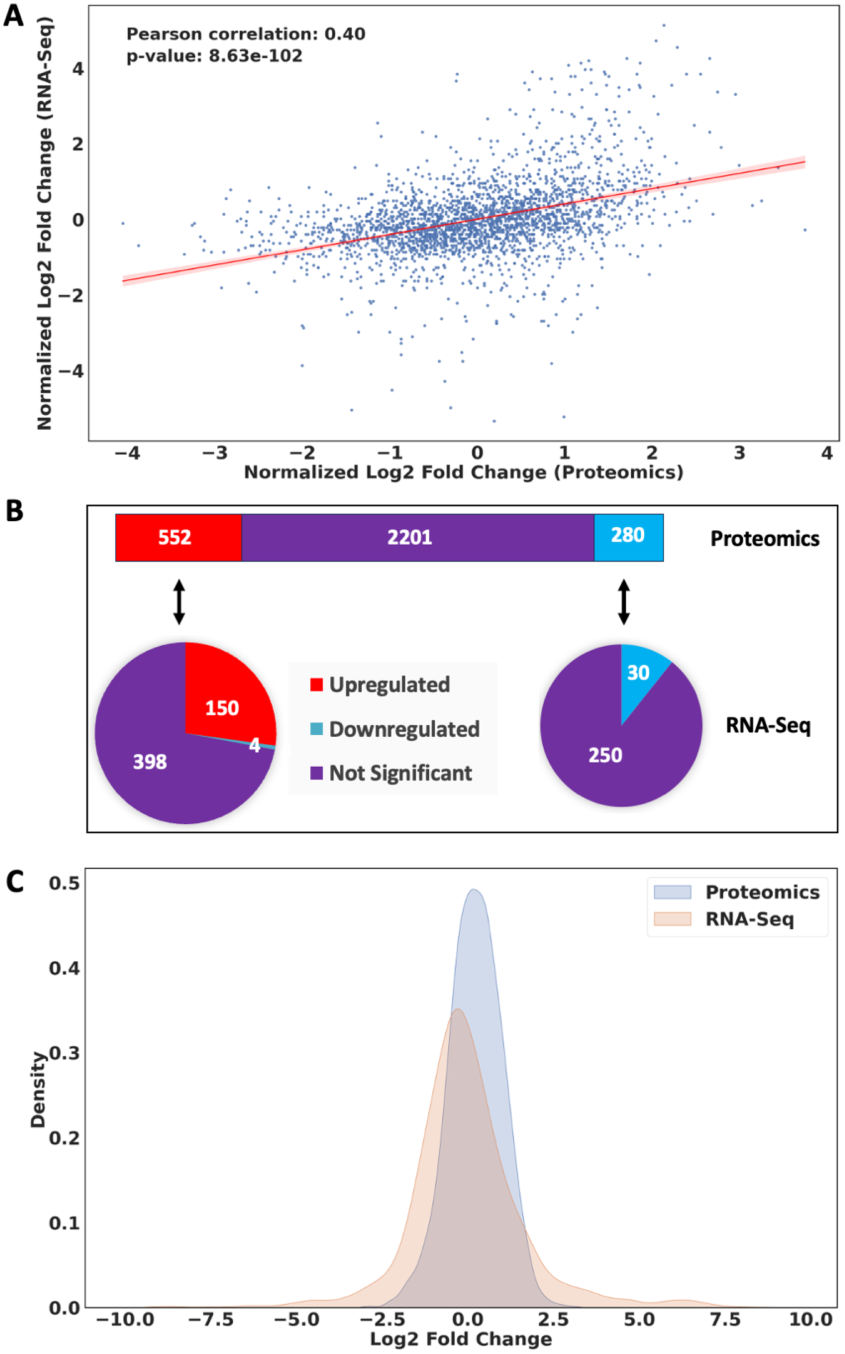
Correlation of RNA-seq and quantitative proteomics data. A. Correlation between proteomics and RNA-seq data, with a Pearson correlation coefficient of 0.4 and a p-value of 8.63 x 10^-102^. The x-axis represents the normalized log2-fold change in proteomics, while the y-axis represents the normalized log2 fold change in RNA-seq data. Each point represents a protein/gene pair, and the red fitted regression line illustrates the trend. B. The distribution of differentially expressed proteins and their corresponding gene expression changes. The upper bar diagram shows downregulated (blue) and upregulated (red) proteins. The pie charts reflect the proportion of upregulated (left) and downregulated (right) proteins that are upregulated (red) or downregulated (blue) in the RNA-seq data. C. Density plot of log2-fold changes from proteomics (blue) and RNA- seq (orange) data. The x-axis represents the log2-fold change, and the y-axis represents the density of the data points.

We also compared the number of differentially expressed genes and proteins. From the 552 upregulated proteins identified in the proteomics data, 150 were also upregulated in the RNA-Seq data, while 4 were downregulated and 398 were not significant (given the applied cutoffs). Similarly, of the 280 downregulated proteins, 30 were also downregulated in the RNA-Seq data, while 250 were not significant (Figure 6B). Overall, we found that 180 proteins had the same expression trend in both datasets, whereas only 4 proteins exhibited inverse regulation (Supplemental Table S6). Two pathways were upregulated in both the RNA-Seq and the proteomics data, benzoate degradation and O-antigen nucleotide sugar biosynthesis.

The distribution of log2-fold changes was compared between RNA-Seq and proteomics data using a density plot (Figure 6C). The distribution of log2-fold changes in RNA-Seq data was broader and of lower amplitude, while the proteomics data showed a narrower and taller distribution (the corresponding distributions are visualized by a violin plot in Supplemental Figure S8). This indicates that the wider range of expression changes seen at the mRNA levels is not reflected at the protein level. This difference speaks to marked post-transcriptional regulation.

### Regulation of RpoS

RpoS is thought to have arisen from an RpoD duplication event prior to the emergence of Proteobacteria and is conserved in proteobacterial species, however, RpoS regulons are not well conserved between species.^55^ While RpoS levels are low in exponential phase, they increase markedly in stationary phase due to a combination of regulatory events.^7^ Such pattern was not observed in *B. thailandensis*. In the RNA-seq dataset, *rpoS* mRNA levels did not meet the filtration criteria for significant differential expression, with a log2-fold change of only 1.2. Furthermore, proteomics data also indicated that RpoS was not markedly increased (log2-fold change 1.08). To further investigate these findings, we performed RT-qPCR analysis to examine the temporal expression of *rpoS*. A transient, but modest increase in RpoS expression was observed during the initial entry into the stationary phase, followed by a return to initial levels (Figure 7A). By comparison, using *lacZ* fused to the *B. pseudomallei rpoS* promoter, a significant increase was observed on entry into stationary phase, followed by a modest decrease.^56^

**Figure 7.**
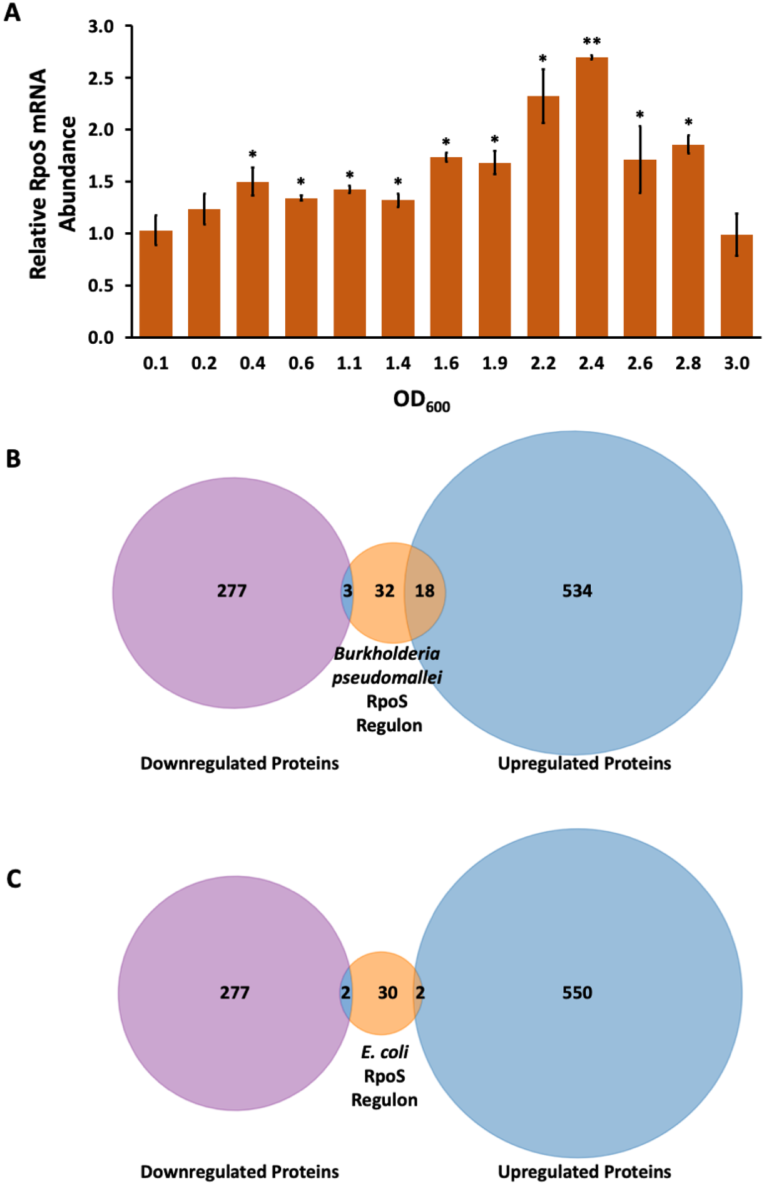
Regulation of RpoS. A. RT-qPCR analysis of *rpoS* gene expression. Relative mRNA abundance is shown as a function of OD_600_, normalized for cell count. Error bars represent the standard deviation from three experiments. Asterisks reflect statistically significant differences compared to the mRNA abundance at OD_600_=0.1 based on a Student’s t-test; *, p<0.05; **, p<0.001. B. Venn diagram comparing differentially expressed proteins regulated by RpoS in *B. pseudomallei* (orange) with differentially expressed proteins in *B. thailandensis* stationary phase; upregulated proteins are shown in blue, and downregulated proteins in purple. C. Venn diagram illustrating the intersection of differentially expressed proteins regulated by RpoS in *E. coli* (orange) with differentially expressed proteins in *B. thailandensis* stationary phase; upregulated proteins are shown in blue, and downregulated proteins in purple.

We examined the RpoS regulon in *Burkholderia pseudomallei*, determined based on mass spectrometric identification of proteins accumulating in an *rpoS* mutant strain.^37^ By comparison to proteins differentially expressed in *B. thailandensis* stationary phase, we identified an only modest overlap (Figure 7B). For example, proteins such as polyphosphate kinase 2 (Ppk2), carboxymuconolactone decarboxylase (PcaC), phasin (PhaP) and a nonribosomally encoded peptide/polyketide synthase (CmaB) were upregulated in *B. thailandensis* during the stationary phase, and they were part of the *B. pseudomallei* RpoS regulon (Supplemental Table S7).

We also compared our findings with the *E. coli* RpoS regulon, likewise identified based on identification of accumulating proteins in an *rpoS* mutant strain.^38^ The overlap was minimal, suggesting distinct regulatory mechanisms between these species (Figure 7C). For instance, while *E. coli* RpoS positively regulates the gene encoding GroES (10 kDa chaperonin), it was downregulated in *B. thailandensis* stationary phase (Supplemental Table S8). Taken together, our findings suggest that *B. thailandensis* employs unique regulatory mechanisms for RpoS that do not result in its marked accumulation during stationary phase.

## Discussion

The transition to the stationary phase represents a crucial adaptive strategy that enables bacteria to endure environmental stress, nutrient scarcity, and other harsh conditions. Such adverse conditions are likely the norm rather than the exception, and they may for instance characterize a host environment rife with antibacterial defenses. For these reasons, the physiology and metabolism of stationary phase cells has been characterized in many species, leading to the identification of several general features. Consistent with such expectations, we found that *B. thailandensis* undergoes extensive metabolic reprogramming, including a shift toward energy conservation, the activation of alternative metabolic pathways, and reduced ribosome biogenesis.^6^ However, our analyses also uncovered unusual regulatory adaptations, for instance involving the sigma factor RpoS.

Integrating transcriptomic and proteomic data has the advantage of providing insights into the molecular mechanisms that underlie the transition to stationary phase, beyond the proteome alterations that arise on account of changes in mRNA abundance. As previously reported,^12^ we also found an only limited correspondence between mRNA and protein abundance (Figure 6). Such modest correlation speaks to post-transcriptional regulation playing a marked role in determining protein abundance. These events may include regulation at the level of translation, perhaps with a contribution from small non-coding RNAs (sRNAs).^57^ In *Burkholderia cepacia* for instance, a number of sRNAs have been characterized and shown to interact with target mRNAs, assisted by RNA chaperones such as Hfq; such interactions may for example lead to occlusion of the ribosome binding site and impaired initiation of translation.^58^ Protein stability may also be regulated, perhaps directed by post-translational modifications.^59^

### Energy-Saving Measures

*B. thailandensis* significantly downregulates nitrogen metabolism, motility, and ribosome biogenesis during the stationary phase, mirroring similar adaptations in other bacteria. Our proteomic analysis identified a marked reduction in core ribosomal proteins and elongation factors, reinforcing the tenet that minimizing ribosome biogenesis is a universal strategy among bacteria during nutrient-limited conditions.^9^

Additionally, the repression of motility-related pathways, such as flagellar assembly and chemotaxis, reflects the expected strategic shift from a motile to a sessile lifestyle, prioritizing biofilm formation as a survival mechanism under nutrient stress.^41^ This energy-saving response is observed across various species as flagellar gene repression reduces the energetic cost of motility. Several gamma-proteobacterial species have even been shown to actively disassemble their flagellar filaments when nutrients are scarce, effectively conserving energy. This programmed loss of flagella leaves behind only what was termed a “relic” of the ejected flagellar motor, associated with a protein complex thought to prevent periplasmic leakage.^60^

Consistent with the goal of reducing the energetic cost of motility, we observed reduced expression of *fliA*, encoding the flagellar sigma factor required for transcription of flagellar genes. In *E. coli* and related bacteria, *flhDC*, which encodes the transcriptional master regulator FlhDC is regulated at several levels; FlhDC in turn stimulates transcription of numerous genes, including *fliA.*^61^ We observe a modest reduction of *flhDC* mRNA, consistent with a comparable regulatory mechanism in *B. thailandensis* (FlhDC was not confidently detected in the proteome analysis). FlhDC was also reported to regulate flagellar genes in the rice pathogen *Burkholderia glumae.*^62^ Since FliA has also been implicated in driving transcription of chemotaxis-related genes, the reduced accumulation of *fliA* mRNA (and the associated 0.6-fold protein abundance in stationary phase; Supplemental Table S3) likely contributes to the observed repression of motility and chemotaxis.

### Metabolic Changes

In the context of nitrogen metabolism, we observed differential expression of two nitrate reductase operons. *BTH_I1852-1854* is expected to be under control of nitrate and nitrite, and this operon was among the most downregulated in stationary phase, whereas *BTH_II1249-1251* was upregulated (Table 1 and Supplemental Figure S2). While regulation of nitrogen metabolism in stationary phase is variable depending on both species and specific environments, upregulation of a membrane-bound nitrate reductase was seen in *Pseudomonas aeruginosa* exposed to the low oxygen environment of the cystic fibrosis lung.^63^ This would be equivalent to the oxygen limitation experienced during stationary phase due to high cell density and limited diffusion. Since nitrate may be used as an alternate electron acceptor when oxygen is limiting, this might lead to its depletion in stationary phase, rationalizing the downregulation of the nitrate reductase encoded by *BTH_I1852-1854*.

Interestingly, *B. thailandensis* upregulates pathways responsible for degradation of the aromatic carbon sources benzoate and anthranilate (Table 1 and Supplemental Figure S3). Both compounds are converted to catechol,^47^ which enters the β-ketoadipate pathway and is converted to intermediates in the citric acid cycle, suggesting a strategic redirection of metabolic flux toward alternative carbon sources when preferred nutrients are limited. In *P. aeruginosa*, anthranilate has been shown to accumulate transiently in stationary phase, and this leads to induction of the operon encoding the dioxygenase, which converts anthranilate to catechol.^64^ Anthranilate is a pathway intermediate in the catabolism of tryptophan,^65^ which is also upregulated in stationary phase (Figure 4A and Supplemental Table S4), indicating a possible source of this metabolite. Notably, anthranilate was shown to regulate pathogenicity-related phenotypes in *P. aeruginosa* such as biofilm formation and antibiotic sensitivity, suggesting a need for careful regulation of its cellular levels.^64^ We also note that anthranilate is an intermediate in the synthesis of 4-hydroxy-3-methyl-2-alkenylquinolines (HMAQ), which are analogous to the *Pseudomonas* quinolone signal PQS, although they do not appear to have comparable roles in quorum sensing;^66^ the increase in anthranilate catabolism correlates with downregulation of genes involved in biosynthesis of HMAQ (Supplemental Table S1).

The observed upregulation of alternative energy pathways such as fatty acid degradation further exemplifies the metabolic flexibility of *B. thailandensis*. This adaptive strategy is reminiscent of metabolic shifts seen in other bacteria.^10^ Fatty acids released from degradation of membrane lipids are scavenged and converted to acyl-CoA, which may then be degraded by the process of β-oxidation. Phosphate acetyl/butyryltransferase was predicted as a critical hub in the PPI network, linked to acetyl-CoA production, the glyoxylate cycle, and fatty acid degradation. Its interactions with proteins involved in polyhydroxyalkanoate (PHA) metabolism further highlight the importance of energy storage and mobilization during the stationary phase.

Our findings also indicate a significant induction of glutathione metabolism, reflecting the expected response to oxidative stress during the stationary phase. The upregulation of glutathione S-transferase and related enzymes suggests a targeted strategy to mitigate reactive oxygen species (ROS), a common challenge under nutrient limitation.^67^

The increase in ROS during stationary phase may also impact iron-sulfur cluster proteins, which are used for a range of essential cellular functions. These clusters are very sensitive to oxidative stress, which can lead to their destabilization or degradation. The stress-responsive Suf (sulfur mobilization) and the house-keeping Isc (iron-sulfur cluster) systems are responsible for iron-sulfur cluster biogenesis and repair.^68^ In *E. coli*, expression of the Isc machinery is regulated by iron availability, but not by oxidative stress, although oxidative stress was shown to inactivate the Isc system.^69^ That Isc protein levels are maintained despite this inactivation was inferred to suggest an alternate function. By contrast, *suf* genes are upregulated in response to oxidative stress. We observed a significant depletion of the IscSUA proteins in stationary phase (Table 2), a depletion that to our knowledge has not been previously reported in other bacterial species. Curiously, expression of the *iscSUA* operon was increased ∼2-fold (Supplemental Table S1), indicating regulation of protein abundance post-transcriptionally. By contrast, *suf* genes were significantly repressed (Supplemental Table S1), although proteins abundance was only modestly reduced (Supplemental Table S3). Our data suggest that the metabolic priorities in stationary phase include a reduced need for iron-sulfur cluster biogenesis.

### RpoS

One of the striking findings of this analysis is that RpoS levels only modestly increased during stationary phase. RpoS is only found in Proteobacteria, and genes have likely been recruited to the RpoS regulons through different selective pressures. As a consequence, RpoS regulons vary between species.^7, 55^ In *E. coli*, the cellular concentration of RpoS was found to increase several fold in stationary phase due to a combination of very complex regulatory events.^70^ Such increase favors RpoS as it competes with the housekeeping sigma factor, σ^70^, for binding to RNA polymerase.^71^ Some of these regulatory events, including transcriptional control and the function of proteases in RpoS degradation during exponential phase were also reported in *P. aeruginosa*.^72^ In other bacterial species, including *B. pseudomallei*, a marked increase in *rpoS* expression has likewise been reported on entry into stationary phase.^56^ The apparent divergence from the canonical regulation of RpoS observed in species such as *E. coli* and *B. pseudomallei* is intriguing and suggests a possible evolution of RpoS-independent regulatory networks in *B. thailandensis* and/or RpoS-mediated regulation of gene expression that is not dependent on marked changes in protein levels.

RpoS bypass mechanisms are not without precedent, and some bacteria even lack RpoS homologs. For instance, the foodborne pathogen *Campylobacter jejuni* lacks stress-related sigma factors such as RpoS, and it relies instead on RpoN (σ^54^).^73^ In other species, such as *Vibrio cholerae*, sigma factor RpoE is important for adaptation to stationary phase along with RpoS, such that RpoS is essential for a general stress response whereas RpoE is more specialized and crucial for responding to envelope stress and maintaining cell viability.^74^ In *B. pseudomallei*, RpoE has been associated with regulation of genes linked to processes such as stress responses and metabolic pathways,^75^ suggesting that *B. thailandensis* RpoE may likewise participate in effecting differential gene expression in stationary phase. RpoN, named for its role in nitrogen assimilation and metabolism in *E. coli*, has been implicated in regulation of genes associated with a range of processes, from flagellar motility to O-antigen expression in various bacterial species.^76^ *Burkholderia* species encode two RpoN homologs, which have been linked to intracellular survival and regulation of type III secretion system genes.^77^ These considerations point to a possible role for *B. thailandensis* RpoE and RpoN in differential gene expression in stationary phase. It is also conceivable that secondary messengers such as (p)ppGpp, which accumulates during the stringent response imposed by nutrient limitation, affect the ability of alternate sigma factors such as RpoS to associate with core RNA polymerase, without depending on significant changes in protein levels, thus resulting in differential gene expression patterns.^9^

Moreover, it is plausible that *B. thailandensis* employs post-translational modifications, such as phosphorylation, acetylation, or glycosylation, to modulate the activity of either RpoS, other regulatory proteins, or even RNA polymerase subunits to favor association of core RNA polymerase with alternate sigma factors. For instance, tyrosine kinases have been described in *B. cenocepacia* and reported to be involved in growth under nutrient limitation.^78^ Similarly, loss of O-linked protein glycosylation results in global proteome changes.^79^ Exploring the extent of post-translational modifications in *B. thailandensis* could unveil novel aspects of bacterial stress adaptation.

### Conclusions

Taken together, this analysis provides a detailed picture of the molecular changes that occur in *B. thailandensis* during the transition from exponential to stationary phase. The coordinated downregulation of energy-intensive processes, such as nitrogen metabolism and motility, along with the upregulation of alternative carbon source metabolism and stress response pathways, reflects the strategic resource reallocation to ensure survival under nutrient-depleted conditions. The modest change in RpoS protein abundance suggests that *B. thailandensis* may rely on unique regulatory approaches. Notably, the only modest correlation between changes in mRNA and protein abundance suggests need to investigate post-transcriptional regulation, such as roles for sRNA in translation initiation and functional consequences of post-translational modification.

These insights into metabolic adaptation and regulatory flexibility provide a foundational understanding of microbial physiology and open new perspectives for biotechnological exploitation. Our findings open several avenues for future research, particularly into the alternative regulatory networks in *B. thailandensis* that may bypass traditional RpoS control. Moreover, the metabolic flexibility observed in *B. thailandensis* suggests potential biotechnological applications, especially in bioremediation. The ability to degrade aromatic compounds like could be harnessed to degrade environmental pollutants, making it a promising candidate for cleaning contaminated soils and water systems.

## Data Availability

The mass spectrometry proteomics data have been deposited to the ProteomeXchange Consortium via the PRIDE^80^ partner repository with the dataset identifier PXD056625 and 10.6019/PXD056625.

The RNA-seq data discussed in this publication have been deposited in NCBI’s Gene Expression Omnibus^81^ and are accessible through GEO Series accession number GSE 279483 (https://www.ncbi.nlm.nih.gov/geo/query/acc.cgi?acc=GSE279483).

## The following supporting information is available

Figure S1. KEGG pathways for chemotaxis and flagellar motility.

Figure S2. KEGG pathways for nitrogen metabolism.

Figure S3. KEGG pathway for benzoate degradation.

Figure S4. Mass spectrometry quantitative proteomics workflow.

Figure S5. KEGG pathway for ribosomal proteins.

Figure S6. KEGG pathway for butanoate metabolism.

Figure S7. Protein-protein interaction (PPI) network functional clustering and GO enrichment.

Figure S8. Violin plots for DEGs and DEPs.

Table S1. DEGs (Excel file).

Table S2. KEGG pathway lists based on DEGs (Excel file).

Table S3. DEPs (Excel file).

Table S4. KEGG pathway lists based on DEPs (Excel file).

Table S5. PPI networks (Excel file).

Table S6. Inverse expression pattern of genes and proteins.

Table S7. Overlap between DEPs and *B. pseudomallei* RpoS regulon.

Table S8. Overlap between DEPs and *E. coli* RpoS regulon.

Table S9. RT-qPCR primer sequences.

Detailed procedures for TMT labeling and protein fractionation.

## Acknowledgments

Supported by the National Science Foundation (MCB-2153410 to AG). Quantitative PCR was performed at the LSU Genomics Core Facility. Mass spectrometry was performed at the LSU Mass Spectrometry Facility.

## Author Contributions

Conceptualization, AA-T, AG; investigation, AA-T, FD; visualization, AA-T; software, AA-T; formal analysis, AA-T; writing-original draft, AA-T; writing-review and editing, AG; funding acquisition, AG.

## Abbreviations

DEGs: Differentially Expressed Genes
DEPs: Differentially Expressed Proteins
FDR: False Discovery Rate
GO: Gene Ontology
KEGG: Kyoto Encyclopedia of Genes and Genomes
NES: Normalized Enrichment Score
PPI: Protein-Protein Interaction
qPCR: Quantitative Polymerase Chain Reaction
RNA-Seq: RNA Sequencing
TMT: Tandem Mass Tag

## References

(1) Elshafie, H. S.; Camele, I. An overview of metabolic activity, beneficial and pathogenic aspects of Burkholderia Spp. Metabolites 2021, 11 (5), 321.

(2) Lewis, E. R.; Torres, A. G. The art of persistence—the secrets to Burkholderia chronic infections. FEMS Pathogens and Disease 2016, 74 (6), ftw070.

(3) Adaikpoh, B. I.; Fernandez, H. N.; Eustáquio, A. S. Biotechnology approaches for natural product discovery, engineering, and production based on Burkholderia bacteria. Current opinion in biotechnology 2022, 77, 102782.

(4) Wiersinga, W. J.; Van der Poll, T.; White, N. J.; Day, N. P.; Peacock, S. J. Melioidosis: insights into the pathogenicity of Burkholderia pseudomallei. Nature Reviews Microbiology 2006, 4 (4), 272–282. Haraga, A.; West, T. E.; Brittnacher, M. J.; Skerrett, S. J.; Miller, S. I. *Burkholderia thailandensis* as a model system for the study of the virulence-associated type III secretion system of *Burkholderia pseudomallei*. Infection and immunity 2008, 76 (11), 5402–5411, Research Support, N.I.H., Extramural Research Support, Non-U.S. Gov’t. DOI: 10.1128/IAI.00626-08.

(5) Maier, R. M.; Pepper, I. L. Bacterial growth. In Environmental microbiology, Elsevier, 2015; pp 37–56.

(6) Jaishankar, J.; Srivastava, P. Molecular basis of stationary phase survival and applications. Frontiers in microbiology 2017, 8, 2000.

(7) Schellhorn, H. E. Function, evolution, and composition of the RpoS regulon in Escherichia coli. Frontiers in microbiology 2020, 11, 560099.

(8) Dworkin, J. Understanding the stringent response: experimental context matters. Mbio 2023, 14 (1), e03404–03422.

(9) Njenga, R.; Boele, J.; Ozturk, Y.; Koch, H. G. Coping with stress: How bacteria fine-tune protein synthesis and protein transport. J Biol Chem 2023, 299 (9), 105163. DOI: 10.1016/j.jbc.2023.105163 From NLM Medline.

(10) Jimenez-Diaz, L.; Caballero, A.; Segura, A. Regulation of Fatty Acids Degradation in Bacteria. In Aerobic Utilization of Hydrocarbons, Oils and Lipids, Rojo, F. Ed.; Springer International Publishing, 2017; pp 1–20.

(11) Craney, A.; Ahmed, S.; Nodwell, J. Towards a new science of secondary metabolism. The Journal of antibiotics 2013, 66 (7), 387–400. Thapa, S. S.; Grove, A. Do Global Regulators Hold the Key to Production of Bacterial Secondary Metabolites? Antibiotics (Basel) 2019, 8 (4), 160. DOI: 10.3390/antibiotics8040160.

(12) Bathke, J.; Konzer, A.; Remes, B.; McIntosh, M.; Klug, G. Comparative analyses of the variation of the transcriptome and proteome of Rhodobacter sphaeroides throughout growth. BMC genomics 2019, 20, 1–13.

(13) Andrews, S. FastQC: a quality control tool for high throughput sequence data. Cambridge, United Kingdom: 2010.

(14) Krueger, F.; James, F.; Ewels, P.; Afyounian, E.; Weinstein, M.; Schuster-Boeckler, B.; Hulselmans, G.; Sclamons. FelixKrueger/TrimGalore: v0. 6.10-add default decompression path. Zenodo 2023.

(15) Ewels, P.; Magnusson, M.; Lundin, S.; Käller, M. MultiQC: summarize analysis results for multiple tools and samples in a single report. Bioinformatics 2016, 32 (19), 3047–3048.

(16) Winsor, G. L.; Khaira, B.; Van Rossum, T.; Lo, R.; Whiteside, M. D.; Brinkman, F. S. The Burkholderia Genome Database: facilitating flexible queries and comparative analyses. Bioinformatics 2008, 24 (23), 2803–2804.

(17) Kim, D.; Paggi, J. M.; Park, C.; Bennett, C.; Salzberg, S. L. Graph-based genome alignment and genotyping with HISAT2 and HISAT-genotype. Nature biotechnology 2019, 37 (8), 907–915.

(18) Liao, Y.; Smyth, G. K.; Shi, W. featureCounts: an efficient general purpose program for assigning sequence reads to genomic features. Bioinformatics 2014, 30 (7), 923–930.

(19) Love, M. I.; Huber, W.; Anders, S. Moderated estimation of fold change and dispersion for RNA-seq data with DESeq2. Genome biology 2014, 15, 1–21.

(20) Benjamini, Y.; Drai, D.; Elmer, G.; Kafkafi, N.; Golani, I. Controlling the false discovery rate in behavior genetics research. Behavioural brain research 2001, 125 (1-2), 279–284.

(21) Gu, Z.; Eils, R.; Schlesner, M. Complex heatmaps reveal patterns and correlations in multidimensional genomic data. Bioinformatics 2016, 32 (18), 2847–2849.

(22) Inc, P. T. Collaborative data science. Montreal: Plotly Technologies Inc Montral 2015, 376.

(23) Wiśniewski, J. Filter-aided sample preparation: the versatile and efficient method for proteomic analysis. In Methods in enzymology, Vol. 585; Elsevier, 2017; pp 15–27.

(24) Orsburn, B. C. Proteome discoverer—a community enhanced data processing suite for protein informatics. Proteomes 2021, 9 (1), 15.

(25) UniProt: the universal protein knowledgebase in 2023. Nucleic acids research 2023, 51 (D1), D523–D531.

(26) Pedregosa, F. Scikit-learn: Machine learning in python Fabian. Journal of machine learning research 2011, 12, 2825.

(27) Hunter, J. D. Matplotlib: A 2D graphics environment. Computing in science & engineering 2007, 9 (03), 90–95.

(28) Szklarczyk, D.; Gable, A. L.; Nastou, K. C.; Lyon, D.; Kirsch, R.; Pyysalo, S.; Doncheva, N. T.; Legeay, M.; Fang, T.; Bork, P. The STRING database in 2021: customizable protein– protein networks, and functional characterization of user-uploaded gene/measurement sets. Nucleic acids research 2021, 49 (D1), D605–D612.

(29) Hagberg, A.; Swart, P. J.; Schult, D. A. *Exploring network structure, dynamics, and function using NetworkX*; Los Alamos National Laboratory (LANL), Los Alamos, NM (United States), 2008.

(30) De Meo, P.; Ferrara, E.; Fiumara, G.; Provetti, A. Generalized louvain method for community detection in large networks. In 2011 11th international conference on intelligent systems design and applications, 2011; IEEE: pp 88–93.

(31) Kanehisa, M.; Furumichi, M.; Sato, Y.; Kawashima, M.; Ishiguro-Watanabe, M. KEGG for taxonomy-based analysis of pathways and genomes. Nucleic acids research 2023, 51 (D1), D587–D592.

(32) Korotkevich, G.; Sukhov, V.; Budin, N.; Shpak, B.; Artyomov, M. N.; Sergushichev, A. Fast gene set enrichment analysis. biorxiv 2016, 060012.

(33) Wickham, H. Data analysis; Springer, 2016.

(34) Schober, P.; Boer, C.; Schwarte, L. A. Correlation coefficients: appropriate use and interpretation. Anesthesia & analgesia 2018, 126 (5), 1763–1768.

(35) Virtanen, P.; Gommers, R.; Oliphant, T. E.; Haberland, M.; Reddy, T.; Cournapeau, D.; Burovski, E.; Peterson, P.; Weckesser, W.; Bright, J. SciPy 1.0: fundamental algorithms for scientific computing in Python. Nature methods 2020, 17 (3), 261–272.

(36) Waskom, M. L. Seaborn: statistical data visualization. Journal of Open Source Software 2021, 6 (60), 3021.

(37) Osiriphun, Y.; Wongtrakoongate, P.; Sanongkiet, S.; Suriyaphol, P.; Thongboonkerd, V.; Tungpradabkul, S. Identification and characterization of RpoS regulon and RpoS-dependent promoters in Burkholderia pseudomallei. Journal of Proteome Research 2009, 8 (6), 3118–3131.

(38) Collet, A.; Cosette, P.; Beloin, C.; Ghigo, J.-M.; Rihouey, C.; Lerouge, P.; Junter, G.-A.; Jouenne, T. Impact of rpoS deletion on the proteome of Escherichia coli grown planktonically and as biofilm. Journal of proteome research 2008, 7 (11), 4659–4669.

(39) Altschul, S. F.; Gish, W.; Miller, W.; Myers, E. W.; Lipman, D. J. Basic local alignment search tool. Journal of molecular biology 1990, 215 (3), 403–410.

(40) Colin, R.; Ni, B.; Laganenka, L.; Sourjik, V. Multiple functions of flagellar motility and chemotaxis in bacterial physiology. FEMS Microbiol. Rev. 2021, 45 (6). DOI: 10.1093/femsre/fuab038 From NLM Medline.

(41) Pesavento, C.; Becker, G.; Sommerfeldt, N.; Possling, A.; Tschowri, N.; Mehlis, A.; Hengge, R. Inverse regulatory coordination of motility and curli-mediated adhesion in *Escherichia coli*. Genes Dev. 2008, 22 (17), 2434–2446. DOI: 10.1101/gad.475808 From NLM Medline.

(42) Chang, L.; Wei, L. I.; Audia, J. P.; Morton, R. A.; Schellhorn, H. E. Expression of the *Escherichia coli* NRZ nitrate reductase is highly growth phase dependent and is controlled by RpoS, the alternative vegetative sigma factor. Mol Microbiol 1999, 34 (4), 756–766. DOI: 10.1046/j.1365-2958.1999.01637.x From NLM Medline.

(43) Glasser, N. R.; Kern, S. E.; Newman, D. K. Phenazine redox cycling enhances anaerobic survival in *Pseudomonas aeruginosa* by facilitating generation of ATP and a proton-motive force. Mol Microbiol 2014, 92 (2), 399–412. DOI: 10.1111/mmi.12566.

(44) Blahut, M.; Sanchez, E.; Fisher, C. E.; Outten, F. W. Fe-S cluster biogenesis by the bacterial Suf pathway. Biochim Biophys Acta Mol Cell Res 2020, 1867 (11), 118829. DOI: 10.1016/j.bbamcr.2020.118829 From NLM Medline.

(45) Diaz, E.; Jimenez, J. I.; Nogales, J. Aerobic degradation of aromatic compounds. Curr. Opin. Biotechnol. 2013, 24 (3), 431–442. DOI: 10.1016/j.copbio.2012.10.010.

(46) Chang, H. K.; Mohseni, P.; Zylstra, G. J. Characterization and regulation of the genes for a novel anthranilate 1,2-dioxygenase from *Burkholderia cepacia* DBO1. J. Bacteriol. 2003, 185 (19), 5871–5881. DOI: 10.1128/JB.185.19.5871-5881.2003 From NLM Medline.

(47) Yanofsky, C. The enzymatic conversion of anthranilic acid to indole. J Biol Chem 1956, 223 (1), 171–184. From NLM Medline.

(48) Puchalska, P.; Crawford, P. A. Multi-dimensional Roles of Ketone Bodies in Fuel Metabolism, Signaling, and Therapeutics. Cell Metab 2017, 25 (2), 262–284. DOI: 10.1016/j.cmet.2016.12.022 From NLM Medline.

(49) Masip, L.; Veeravalli, K.; Georgiou, G. The many faces of glutathione in bacteria. Antioxid Redox Signal 2006, 8 (5-6), 753–762. DOI: 10.1089/ars.2006.8.753 From NLM Medline.

(50) Huang, H.; Grove, A. The transcriptional regulator TamR from *Streptomyces coelicolor* controls a key step in central metabolism during oxidative stress. Mol Microbiol 2013, 87 (6), 1151–1166, Research Support, U.S. Gov’t, Non-P.H.S. DOI: 10.1111/mmi.12156.

(51) Sivapragasam, S.; Grove, A. The Link between Purine Metabolism and Production of Antibiotics in *Streptomyces*. Antibiotics (Basel) 2019, 8 (2), 76. DOI: 10.3390/antibiotics8020076.

(52) Wang, Z.; Xie, X.; Shang, D.; Xie, L.; Hua, Y.; Song, L.; Yang, Y.; Wang, Y.; Shen, X.; Zhang, L. A c-di-GMP Signaling Cascade Controls Motility, Biofilm Formation, and Virulence in *Burkholderia thailandensis*. Appl. Environ. Microbiol. 2022, 88 (7), e0252921. DOI: 10.1128/aem.02529-21 From NLM Medline.

(53) Schoelmerich, M. C.; Müller, V. Energy-converting hydrogenases: the link between H2 metabolism and energy conservation. Cellular and Molecular Life Sciences 2020, 77 (8), 1461–1481.

(54) Picard, F.; Dressaire, C.; Girbal, L.; Cocaign-Bousquet, M. Examination of post-transcriptional regulations in prokaryotes by integrative biology. Comptes rendus biologies 2009, 332 (11), 958–973. Zhang, D.; Li, S. H.-J.; King, C. G.; Wingreen, N. S.; Gitai, Z.; Li, Z. Global and gene-specific translational regulation in Escherichia coli across different conditions. PLoS Comput. Biol. 2022, 18 (10), e1010641.

(55) Chiang, S. M.; Schellhorn, H. E. Evolution of the RpoS regulon: origin of RpoS and the conservation of RpoS-dependent regulation in bacteria. J. Mol. Evol. 2010, 70 (6), 557–571. DOI: 10.1007/s00239-010-9352-0 From NLM Medline.

(56) Subsin, B.; Thomas, M. S.; Katzenmeier, G.; Shaw, J. G.; Tungpradabkul, S.; Kunakorn, M. Role of the stationary growth phase sigma factor RpoS of *Burkholderia pseudomallei* in response to physiological stress conditions. J. Bacteriol. 2003, 185 (23), 7008–7014. DOI: 10.1128/JB.185.23.7008-7014.2003 From NLM Medline.

(57) Papenfort, K.; Melamed, S. Small RNAs, Large Networks: Posttranscriptional Regulons in Gram-Negative Bacteria. Annu. Rev. Microbiol. 2023, 77, 23–43. DOI: 10.1146/annurev-micro-041320-025836 From NLM Medline.

(58) Matos, G. R.; Feliciano, J. R.; Leitao, J. H. Non-coding regulatory sRNAs from bacteria of the *Burkholderia cepacia* complex. Appl. Microbiol. Biotechnol. 2024, 108 (1), 280. DOI: 10.1007/s00253-024-13121-6 From NLM Medline.

(59) Macek, B.; Forchhammer, K.; Hardouin, J.; Weber-Ban, E.; Grangeasse, C.; Mijakovic, I. Protein post-translational modifications in bacteria. Nat Rev Microbiol 2019, 17 (11), 651–664. DOI: 10.1038/s41579-019-0243-0.

(60) Ferreira, J. L.; Gao, F. Z.; Rossmann, F. M.; Nans, A.; Brenzinger, S.; Hosseini, R.; Wilson, A.; Briegel, A.; Thormann, K. M.; Rosenthal, P. B. γ-proteobacteria eject their polar flagella under nutrient depletion, retaining flagellar motor relic structures. PLoS Biology 2019, 17 (3), e3000165.

(61) Fitzgerald, D. M.; Bonocora, R. P.; Wade, J. T. Comprehensive mapping of the *Escherichia coli* flagellar regulatory network. PLoS Genet 2014, 10 (10), e1004649. DOI: 10.1371/journal.pgen.1004649 From NLM Medline.

(62) Kim, J.; Kang, Y.; Choi, O.; Jeong, Y.; Jeong, J. E.; Lim, J. Y.; Kim, M.; Moon, J. S.; Suga, H.; Hwang, I. Regulation of polar flagellum genes is mediated by quorum sensing and FlhDC in *Burkholderia glumae*. Mol Microbiol 2007, 64 (1), 165–179. DOI: 10.1111/j.1365-2958.2007.05646.x From NLM Medline.

(63) Palmer, K. L.; Brown, S. A.; Whiteley, M. Membrane-bound nitrate reductase is required for anaerobic growth in cystic fibrosis sputum. J. Bacteriol. 2007, 189 (12), 4449–4455. DOI: 10.1128/JB.00162-07 From NLM Medline.

(64) Hwang, H. J.; Li, X. H.; Kim, S. K.; Lee, J. H. Anthranilate Acts as a Signal to Modulate Biofilm Formation, Virulence, and Antibiotic Tolerance of *Pseudomonas aeruginosa* and Surrounding Bacteria. Microbiol Spectr 2022, 10 (1), e0146321. DOI: 10.1128/spectrum.01463-21 From NLM Medline.

(65) Kurnasov, O.; Jablonski, L.; Polanuyer, B.; Dorrestein, P.; Begley, T.; Osterman, A. Aerobic tryptophan degradation pathway in bacteria: novel kynurenine formamidase. FEMS microbiology letters 2003, 227 (2), 219–227. DOI: 10.1016/S0378-1097(03)00684-0 From NLM Medline.

(66) Coulon, P. M. L.; Zlosnik, J. E. A.; Deziel, E. Presence of the Hmq System and Production of 4-Hydroxy-3-Methyl-2-Alkylquinolines Are Heterogeneously Distributed between *Burkholderia cepacia* Complex Species and More Prevalent among Environmental than Clinical Isolates. Microbiol Spectr 2021, 9 (1), e0012721. DOI: 10.1128/Spectrum.00127-21 From NLM Medline.

(67) Lemire, J.; Alhasawi, A.; Appanna, V. P.; Tharmalingam, S.; Appanna, V. D. Metabolic defence against oxidative stress: the road less travelled so far. J. Appl. Microbiol. 2017, 123 (4), 798–809. DOI: 10.1111/jam.13509 From NLM Medline.

(68) Esquilin-Lebron, K.; Dubrac, S.; Barras, F.; Boyd, J. M. Bacterial Approaches for Assembling Iron-Sulfur Proteins. mBio 2021, 12 (6), e0242521. DOI: 10.1128/mBio.02425-21 From NLM Medline.

(69) Jang, S.; Imlay, J. A. Hydrogen peroxide inactivates the *Escherichia coli* Isc iron-sulphur assembly system, and OxyR induces the Suf system to compensate. Mol Microbiol 2010, 78 (6), 1448–1467. DOI: 10.1111/j.1365-2958.2010.07418.x From NLM Medline.

(70) Lange, R.; Hengge-Aronis, R. The cellular concentration of the sigma S subunit of RNA polymerase in *Escherichia coli* is controlled at the levels of transcription, translation, and protein stability. Genes Dev. 1994, 8 (13), 1600–1612. DOI: 10.1101/gad.8.13.1600 From NLM Medline.

(71) Bouillet, S.; Bauer, T. S.; Gottesman, S. RpoS and the bacterial general stress response. Microbiol. Mol. Biol. Rev. 2024, 88 (1), e0015122. DOI: 10.1128/mmbr.00151-22 From NLM Medline.

(72) Bertani, I.; Sevo, M.; Kojic, M.; Venturi, V. Role of GacA, LasI, RhlI, Ppk, PsrA, Vfr and ClpXP in the regulation of the stationary-phase sigma factor rpoS/RpoS in *Pseudomonas*. Arch. Microbiol. 2003, 180 (4), 264–271. DOI: 10.1007/s00203-003-0586-8 From NLM Medline.

(73) Hwang, S.; Jeon, B.; Yun, J.; Ryu, S. Roles of RpoN in the resistance of *Campylobacter jejuni* under various stress conditions. BMC Microbiol 2011, 11, 207. DOI: 10.1186/1471-2180-11-207 From NLM Medline.

(74) Davis, B. M.; Waldor, M. K. High-throughput sequencing reveals suppressors of *Vibrio cholerae rpoE* mutations: one fewer porin is enough. Nucleic Acids Res 2009, 37 (17), 5757–5767. DOI: 10.1093/nar/gkp568 From NLM Medline.

(75) Thongboonkerd, V.; Vanaporn, M.; Songtawee, N.; Kanlaya, R.; Sinchaikul, S.; Chen, S. T.; Easton, A.; Chu, K.; Bancroft, G. J.; Korbsrisate, S. Altered proteome in *Burkholderia pseudomallei rpoE* operon knockout mutant: insights into mechanisms of *rpoE* operon in stress tolerance, survival, and virulence. J Proteome Res 2007, 6 (4), 1334–1341. DOI: 10.1021/pr060457t From NLM Medline.

(76) Yu, C.; Yang, F.; Xue, D.; Wang, X.; Chen, H. The Regulatory Functions of sigma(54) Factor in Phytopathogenic Bacteria. Int J Mol Sci 2021, 22 (23). DOI: 10.3390/ijms222312692 From NLM Medline.

(77) Diep, D. T. H.; Vong, L. B.; Tungpradabkul, S. Function of *Burkholderia pseudomallei* RpoS and RpoN2 in bacterial invasion, intracellular survival, and multinucleated giant cell formation in mouse macrophage cell line. Antonie Van Leeuwenhoek 2024, 117 (1), 39. DOI: 10.1007/s10482-024-01944-2 From NLM Medline.

(78) Andrade, A.; Tavares-Carreon, F.; Khodai-Kalaki, M.; Valvano, M. A. Tyrosine Phosphorylation and Dephosphorylation in *Burkholderia cenocepacia* Affect Biofilm Formation, Growth under Nutritional Deprivation, and Pathogenicity. Appl. Environ. Microbiol. 2016, 82 (3), 843–856. DOI: 10.1128/AEM.03513-15 From NLM Medline.

(79) Oppy, C. C.; Jebeli, L.; Kuba, M.; Oates, C. V.; Strugnell, R.; Edgington-Mitchell, L. E.; Valvano, M. A.; Hartland, E. L.; Newton, H. J.; Scott, N. E. Loss of O-Linked Protein Glycosylation in *Burkholderia cenocepacia* Impairs Biofilm Formation and Siderophore Activity and Alters Transcriptional Regulators. mSphere 2019, 4 (6). DOI: 10.1128/mSphere.00660-19.

(80) Perez-Riverol, Y.; Bai, J.; Bandla, C.; Garcia-Seisdedos, D.; Hewapathirana, S.; Kamatchinathan, S.; Kundu, D. J.; Prakash, A.; Frericks-Zipper, A.; Eisenacher, M.;, et al. The PRIDE database resources in 2022: a hub for mass spectrometry-based proteomics evidences. Nucleic Acids Res 2022, 50 (D1), D543–D552. DOI: 10.1093/nar/gkab1038 From NLM Medline.

(81) Edgar, R.; Domrachev, M.; Lash, A. E. Gene Expression Omnibus: NCBI gene expression and hybridization array data repository. Nucleic Acids Res 2002, 30 (1), 207–210. DOI: 10.1093/nar/30.1.207 From NLM Medline.

