## Supplemental figures and tables for "Transcriptome-proteome profiling in *Burkholderia thailandensis* during the transition from exponential to stationary phase"

**Contents:**

**Supplemental Figures:**

Figure S1. KEGG pathways for chemotaxis and flagellar motility.
Figure S2. KEGG pathways for nitrogen metabolism.
Figure S3. KEGG pathway for benzoate degradation.
Figure S4. Mass spectrometry quantitative proteomics workflow.
Figure S5. KEGG pathway for ribosomal proteins.
Figure S6. KEGG pathway for butanoate metabolism.
Figure S7. Protein-protein interaction (PPI) network functional clustering and GO enrichment.
Figure S8. Violin plots for DEGs and DEPs.

**Supplemental Tables:**

Table S1. DEGs (Excel file).
Table S2. KEGG pathway lists based on DEGs (Excel file).
Table S3. DEPs (Excel file).
Table S4. KEGG pathway lists based on DEPs (Excel file).
Table S5. PPI networks (Excel file).
Table S6. Inverse expression pattern of genes and proteins.
Table S7. Overlap between DEPs and *B. pseudomallei* RpoS regulon.
Table S8. Overlap between DEPs and *E. coli* RpoS regulon.
Table S9. RT-qPCR primer sequences.

**Detailed procedures for TMT labeling and protein fractionation**

**Supplemental Figures:**

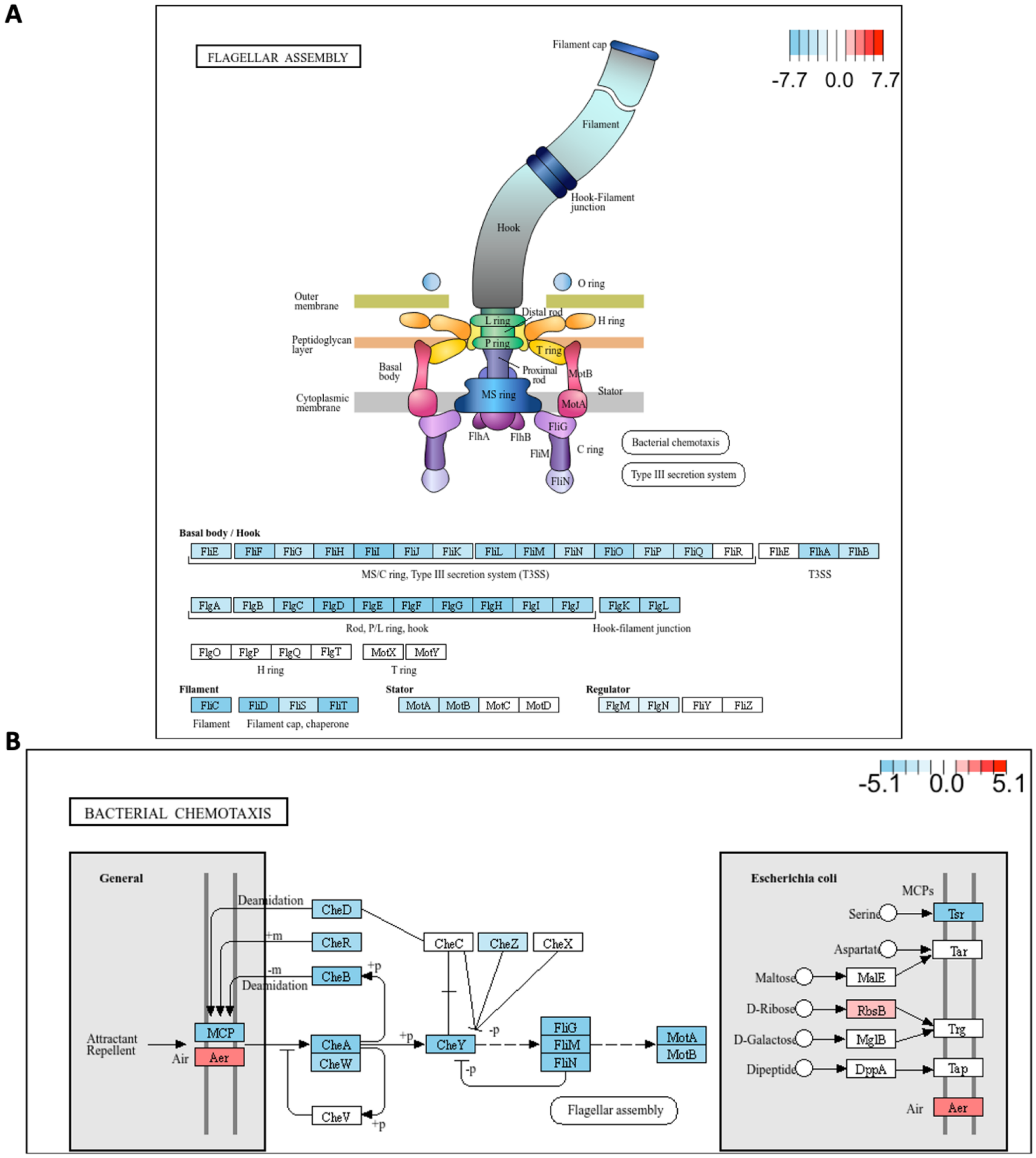


**Figure S1. KEGG pathways for chemotaxis and flagellar motility**. A. Schematic of the bacterial flagellar structure, indicating the arrangement of its main components (top). KEGG pathways for *B. thailandensis* flagellar motility (bottom); see https://www.genome.jp/pathway/bte02040 for identification of locus tags. B. KEGG pathways for bacterial chemotaxis; see https://www.genome.jp/pathway/bte02030 for identification of locus tags. The connection between gene products (rectangles) and chemical compounds (circles) are indicated. DEGs that meet the log2-fold change of |2| are shown in color. The color-coded scales in the top-right corners represent the log2-fold changes in gene expression in stationary phase, with shades of red reflecting upregulation and shades of blue representing downregulation.

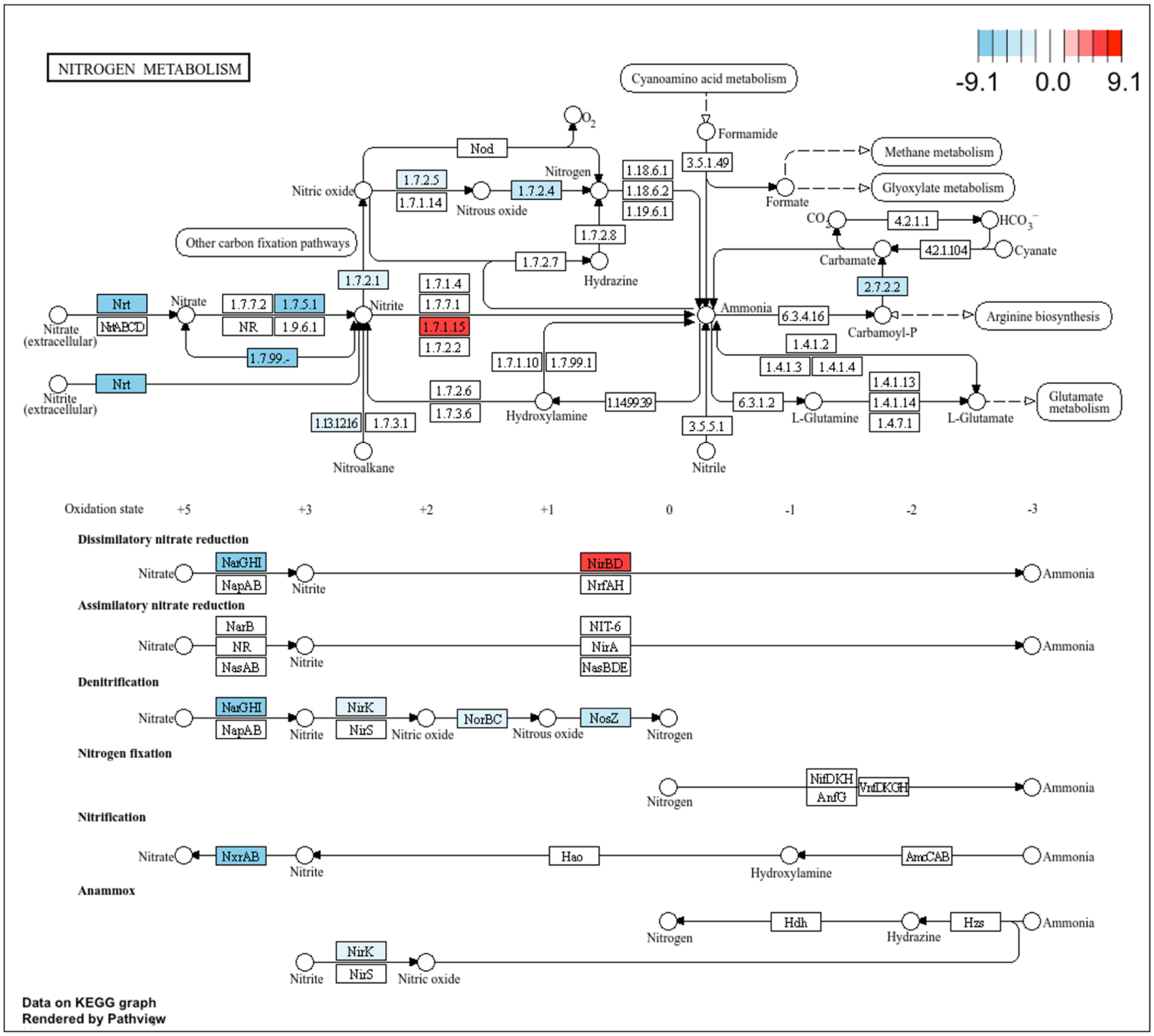


**Figure S2. KEGG pathways for nitrogen metabolism**. KEGG pathways for different aspects of nitrogen metabolism; see https://www.genome.jp/pathway/bte00910 for identification of locus tags. The connection between gene products (rectangles) and chemical compounds (circles) are indicated. DEGs that meet the log2-fold change of |2| are shown in color. The color-coded scale in the top-right corner represents the log2-fold changes in gene expression in stationary phase, with shades of red reflecting upregulation and shades of blue representing downregulation.

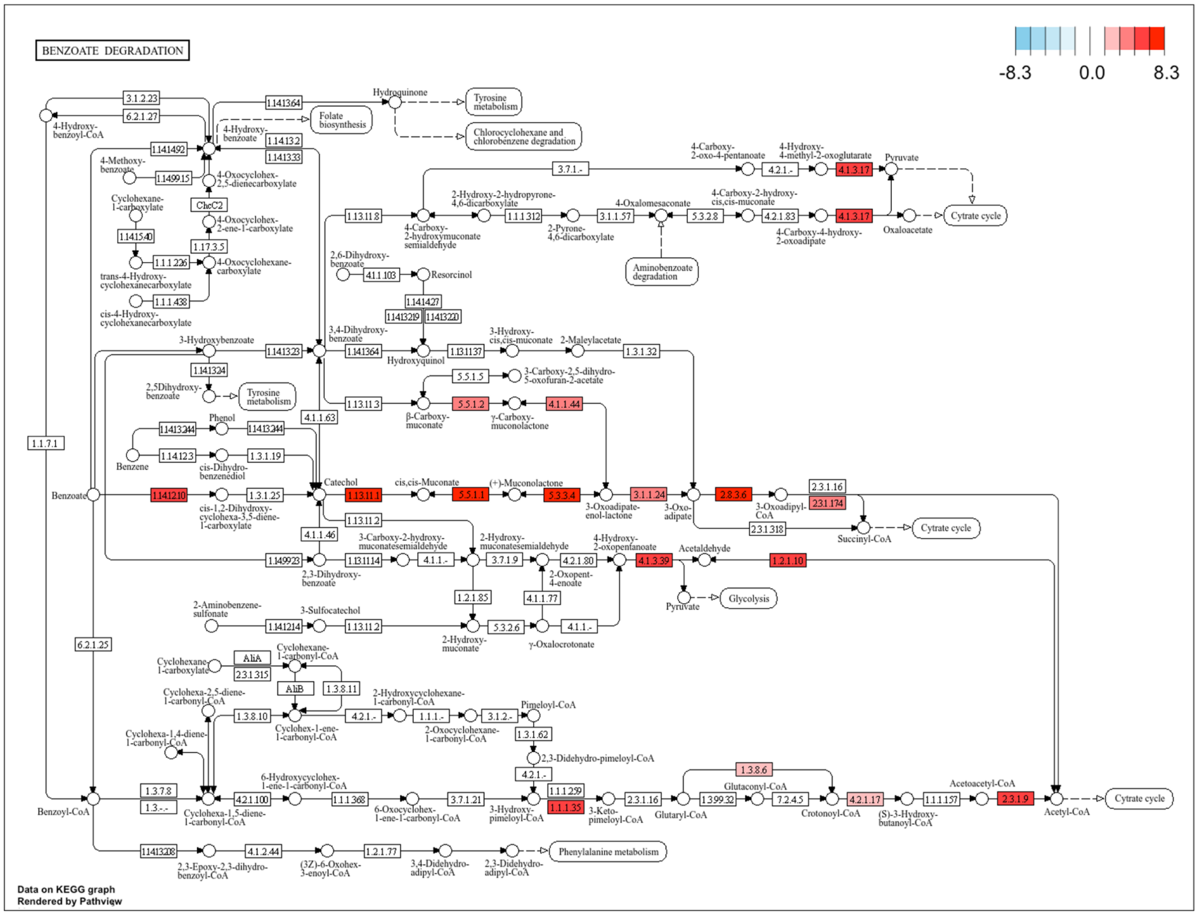


**Figure S3. KEGG pathway for benzoate degradation**. See https://www.genome.jp/pathway/bte00362 for identification of locus tags. The connection between gene products (rectangles) and chemical compounds (circles) are indicated. DEGs that meet the log2-fold change of |2| are shown in color. The color-coded scale in the top-right corner represents the log2-fold changes in gene expression in stationary phase, with shades of red reflecting upregulation and shades of blue representing downregulation.

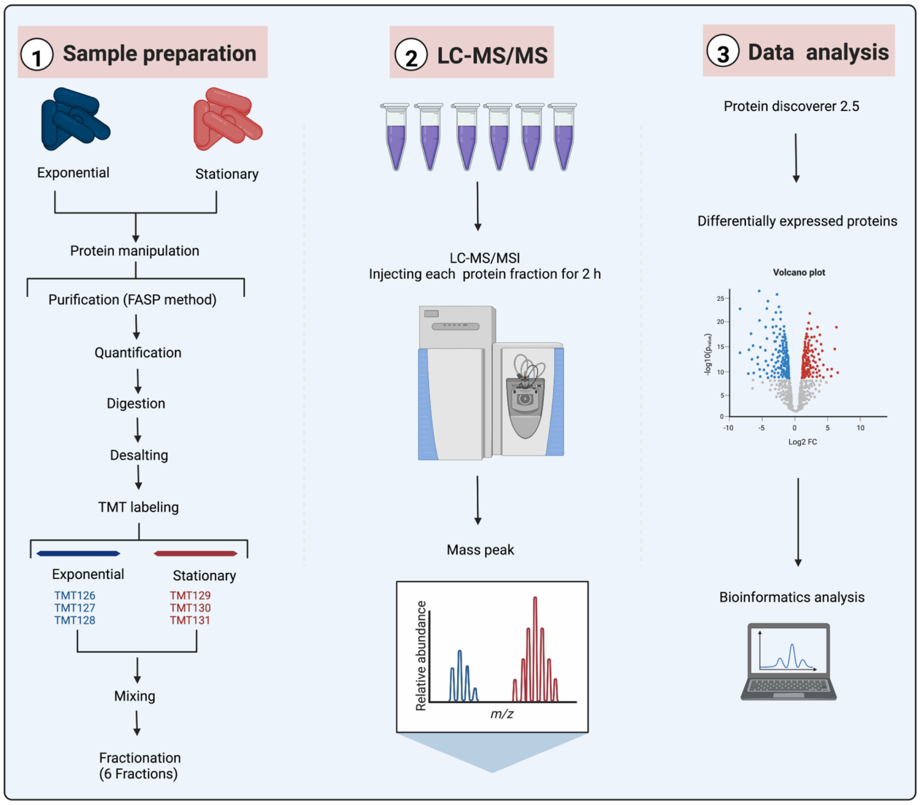


**Figure S4. Mass spectrometry quantitative proteomics workflow**. 1. The samples underwent protein manipulations such as reduction by DTT, followed by purification using the Filter-Aided Sample Preparation (FASP) method, protein quantification, enzymatic digestion, and desalting. The resulting peptides were labeled with Tandem Mass Tags (TMT) to enable multiplexed quantification. TMT126, TMT127, and TMT128 were used for labeling samples from the exponential phase, while TMT129, TMT130, and TMT131 were used for the stationary phase. The labeled samples were then mixed and separated into six fractions. 2. Each fraction was subjected to liquid chromatography-tandem mass spectrometry (LC-MS/MS). Peptides were separated based on their mass-to-charge ratio (m/z) and quantified. 3. The mass spectrometry data were processed using the Protein Discoverer 2.5 software to determine differential accumulation of proteins followed by bioinformatics analysis. Created with BioRender.com.

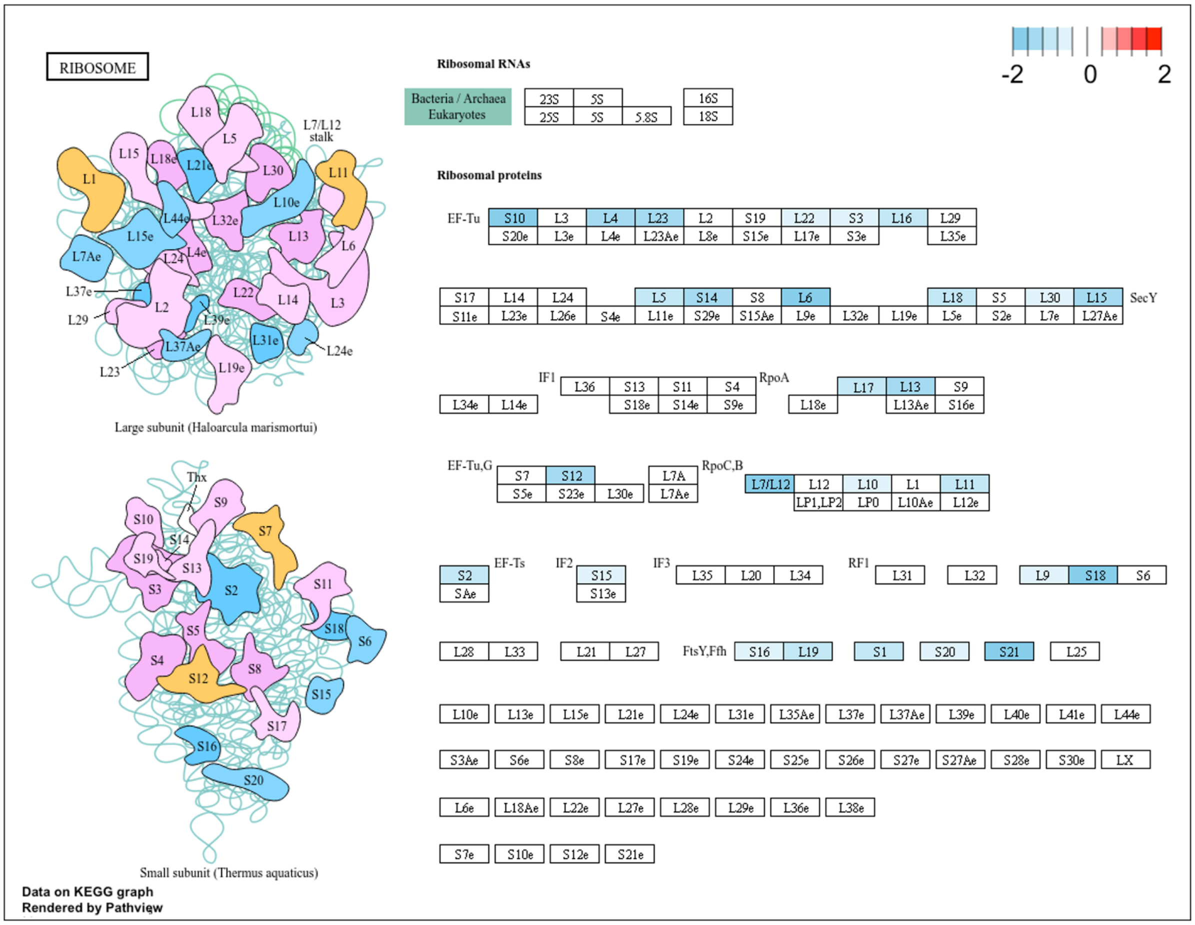


**Figure S5. KEGG pathway for ribosomal proteins**. A schematic representation of the ribosome (left), identifying ribosomal proteins. Ribosomal proteins and translation factors are depicted at the right; see https://www.genome.jp/pathway/bte03010 for identification of locus tags. DEPs that meet the log2-fold change of |0.5| are shown in color. The color-coded scale in the top-right corner represents the log2-fold changes in gene expression in stationary phase, with shades of red reflecting upregulation and shades of blue representing downregulation.

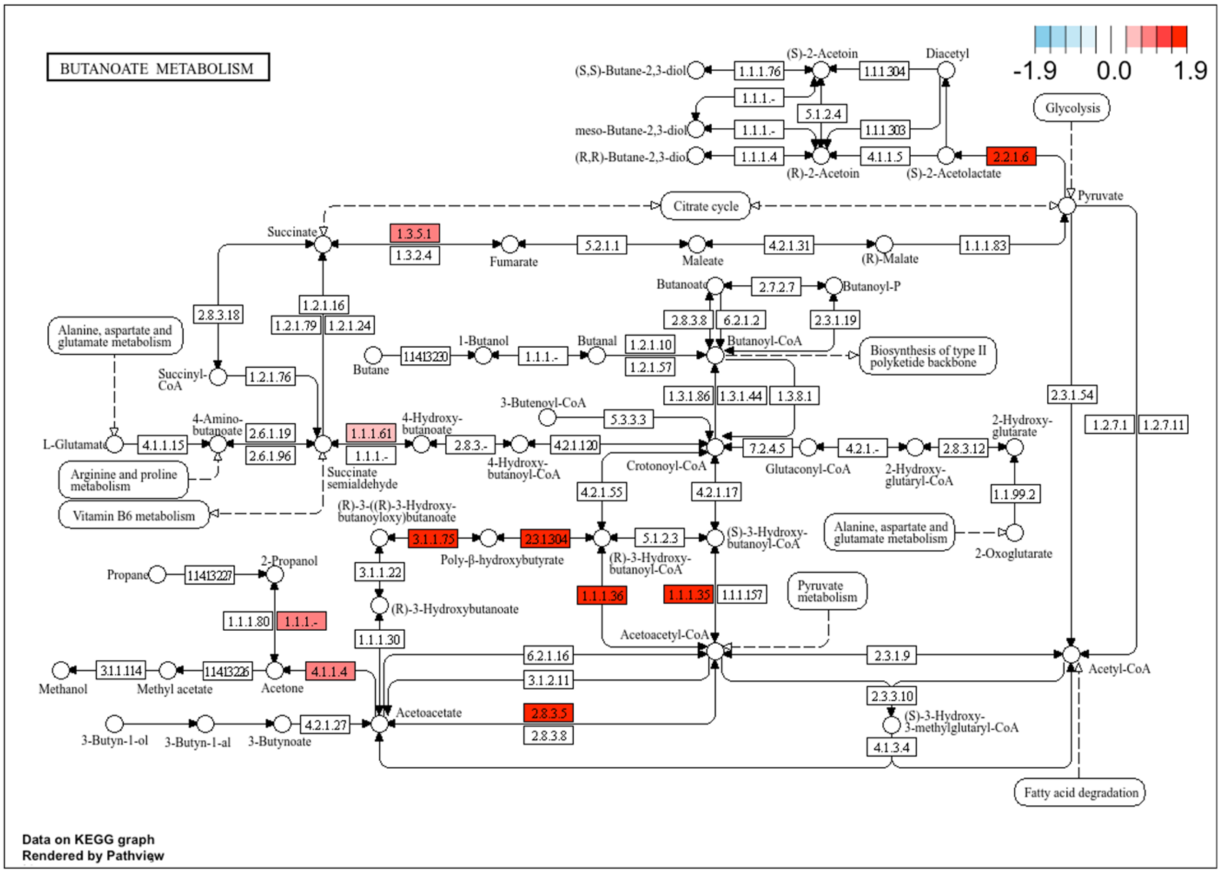


**Figure S6. KEGG pathway for butanoate metabolism**. See https://www.genome.jp/pathway/bte00650 for identification of locus tags. The connection between gene products (rectangles) and chemical compounds (circles) are indicated. DEPs that meet the log2-fold change of |0.5| are shown in color. The color-coded scale in the top-right corner represents the log2-fold changes in gene expression in stationary phase, with shades of red reflecting upregulation and shades of blue representing downregulation.

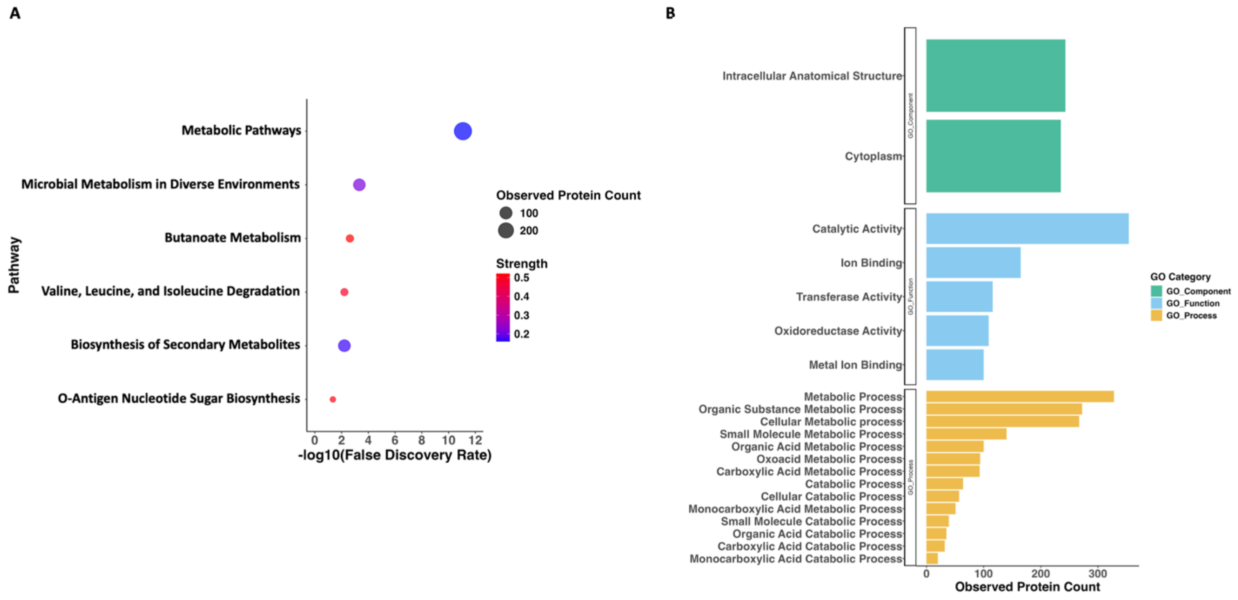


**Figure S7. Protein-protein interaction (PPI) network functional clustering and GO enrichment.** A. Bubble plot identifying the KEGG pathways associated with the PPI network during the stationary phase. The x-axis lists the pathway descriptions, while the y-axis represents the -log10(p-value) of the enrichment. Each bubble represents a pathway, with the size denoting the proportion of proteins in the pathway that are differentially expressed. The color gradient of the bubbles indicates the enrichment strength, with red shades representing stronger enrichment. B. GO enrichment analysis of upregulated proteins within the PPI network. The x-axis indicates the observed gene count associated with each GO term, and the y-axis lists the descriptions of the enriched terms. The plot distinguishes between different GO categories using color: Biological process (yellow), molecular function (blue), and cellular component (green).

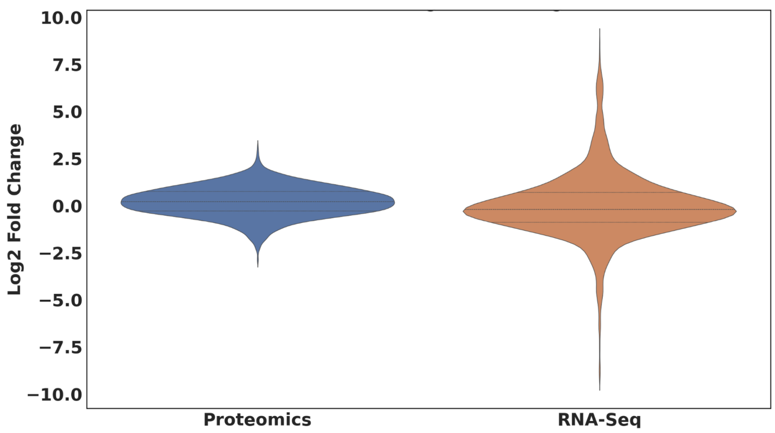


**Figure S8. Violin plots for DEGs and DEPs.** Distribution of log2-fold changes for proteomics (blue) and RNA-Seq (orange) datasets. Each violin plot provides a kernel density estimation of the data. For the proteomics dataset, the median log2-fold change is 0.21, with an interquartile range (IQR) of 1.04. The 95% confidence interval for the mean log2-fold change ranges from 0.19 to 0.25. The RNA-Seq dataset has a median log2-fold change of -0.2, with a broader interquartile range (IQR) of 1.6. The 95% confidence interval for the mean log2-fold change ranges from -0.09 to 0.04.

**Supplemental Tables:**

**Table S6. Inverse expression pattern of genes and proteins.**

| Gene ID | Gene Log2 Fold Change | Corresponding Protein ID | Protein Log2 Fold Change | Name of protein |
|---|---|---|---|---|
| BTH_I2387 | -4.6 | Q2SVZ1 | 0.81 | Oxidoreductase, short-chain dehydrogenase/reductase family |
| BTH_II1074 | -2.3 | Q2T6C8 | 0.50 | HTH-type transcriptional regulator BetI |
| BTH_II0445 | -2.2 | Q2T854 | 0.95 | ABC transporter, ATP-binding protein |
| BTH_I3168 | -6.5 | Q2STT7 | 0.88 | Flagellar biosynthesis protein FlhF |

**Table S6**. Inverse regulation in *B. thailandensis* stationary phase of DEGs and DEPs that meet both the log2-fold change of |2| for DEGs and |0.5| for DEPs.

**Table S7. Overlap between DEPs and *B. pseudomallei* RpoS regulon.**

| Functional Annotation | *ΔrpoS* *B. pseudomallei* | Regulation in *B. thailandensis* | Identity (%) |
|---|---|---|---|
| Polyphosphate kinase 2 family (Ppk2) | Down | Up | 96 |
| Carboxymuconolactone decarboxylase (PcaC) | Down | Up | 95 |
| Nonribosomally encoded peptide/polyketide synthase (CmaB) | Down | Up | 97 |
| HSP20/alpha crystallin family protein | Down | Up | 93 |
| Universal stress protein family | Up | Up | 96 |
| Universal stress protein family | Down | Up | 98 |
| Universal stress protein UspA | Down | Up | 91 |
| Phasin (PhaP) | Down | Up | 100 |
| Alkyl hydroperoxide reductase D (AhpD) | Down | Up | 94 |
| Antioxidant, AhpC/Tsa family | Down | Up | 98 |
| Osmotically inducible Y domain protein (OsmY) | Down | Up | 92 |
| Osmotically inducible Y domain protein (OsmY) | Down | Up | 95 |
| NADPH-dependent FMN reductase | Down | Up | 92 |
| Inclusion body family protein | Down | Up | 33 |
| Ribosomal natural product, two-chain TOMM family | Down | Up | 76 |
| DUF1842 domain-containing protein | Down | Up | 85 |
| Hypothetical protein BPSS0213 | Down | Up | 84 |
| Polyketide synthase, putative | Down | Up | 75 |
| 4-hydroxy-3-methylbut-2-enyl diphosphate reductase (IspH) | Up | Down | 99 |
| Chaperonin GroES | Down | Down | 99 |
| Alkyl hydroperoxide reductase C | Down | Down | 31 |

**Table S7**. Proteins for which the corresponding genes were found to be regulated in a *B. pseudomallei ΔrpoS* strain relative to wild-type. Regulation in *B. thailandensis* refers to stationary phase relative to exponential growth; only proteins that meet the log2-fold change of |0.5| are included.

**Table S8. Overlap between DEPs and *E. coli* RpoS regulon.**

| ***ΔrpoS* *E. coli*** | **Protein annotation in *E. coli*** | **Regulation in *B. thailandensis*** | **Protein annotation in *B. thailandensis*** | **Identity (%)** |
|---|---|---|---|---|
| Down | TufA - Elongation factor Tu | Up | Elongation factor Tu (EF-Tu) | 80 |
| Up | AckA - Acetate kinase | Up | Acetate kinase | 36 |
| Up | GroES - 10 kDa chaperonin | Down | Co-chaperonin GroES, 10 kDa chaperonin | 53 |
| Up | FabB – 3-oxoacyl-[acyl-carrier-protein] synthase 1 | Down | 3-oxoacyl-[acyl-carrier-protein] synthase 2 | 38 |

**Table S8**. Proteins for which the corresponding genes were found to be regulated in an *E. coli ΔrpoS* strain relative to wild-type. Regulation in *B. thailandensis* refers to stationary phase relative to exponential growth; only proteins that meet the log2-fold change of |0.5| are included.

**Table S9. RT-qPCR primer sequences.**

| **Name** | **Sequence (5' to 3')** |
|---|---|
| RpoS_FW_BTH_I2226 | ACGACATCGCGTATCTGACC |
| RpoS_RV_BTH_I2226 | GGGTAGAAGATCGAGCAGGC |
| Glutamate synthase_FW_BTH_I3014 | GCAAGAAGAGCCACGAAATC |
| Glutamate synthase_RV_BTH_I3014 | CCATCTCCTCGCGATAGAAC |

**Detailed procedures for TMT labeling and protein fractionation**

Samples were processed using the filter-aided sample preparation (FASP) method,[1] with modifications. Forty mL of the bacterial culture were centrifuged at 17,800 × g for 5 min, followed by two washes with phosphate buffered saline (PBS). The bacterial pellets were stored at -80°C. Protein extraction was carried out using a lysis buffer (8 M urea in 50 mM Tris, pH 8.0). The cell pellets were sonicated twice using a Branson SFX250 Sonifier at 35% output at 10 secs pulse with 1-minute gap in between. Homogenized samples were centrifuged at 8,600 × g for 5 min. Protein concentration was determined using the Pierce™ bicinchoninic acid (BCA) Protein Assay Kit (Thermo Fisher Scientific).

Proteins (100 µg total) were first reduced with 50 mM DTT and then alkylated with iodoacetamide. Subsequent overnight digestion was performed with trypsin at 37°C. The 100 µg quantity was chosen based on the published protocol[1] and past experience. Generally, a loss of up to 50% of the sample is associated with this protocol in a tradeoff for a high peptide sample purity. Accordingly, the generated tryptic digests provided ~50 µg of peptide material for subsequent TMT tagging.

TMTsixplex™ labeling was conducted following the manufacturer's protocol (Thermo Fisher Scientific).

Following labeling, the peptides were pooled and fractionated using strong cation exchange stage tips (AttractSPE Disk, Affinisep, Le Houlme, Normandy, France). These tips require 30 µg of total peptide material. Therefore, we used 5 µg of each TMT reaction for pooling to generate the required 30 µg total for fractionation. Elution was performed following the manufacturer protocol using increasing amounts of ammonium acetate. A total of 6 fractions where dried and resuspended in 0.1% formic acid. The LC-MS/MS analysis was executed on an Orbitrap Fusion Tribrid mass spectrometer using one-half of the reconstituted material.

**References**

(1) Wiśniewski, J. Filter-aided sample preparation: the versatile and efficient method for proteomic analysis. In *Methods in enzymology*, Vol. 585; Elsevier, 2017; pp 15-27.
